# Disrupting the cockroach circadian clock through RNAi-mediated knockdown of clock proteins

**DOI:** 10.64898/2026.04.27.720303

**Authors:** Huleg Zolmon, Tabea Trummel, Lukas Kräling, Patrick Przybylla, Anna C. Schneider, Olaf Stursberg, Monika Stengl

**Author notes:** Both authors contributed equally (Huleg Zolmon, Tabea Trummel). **Corresponding author email address:**, **Corresponding author mailing address:** Anna C. Schneider Universität Kassel, FB 10 Biologie, Tierphysiologie Heinrich-Plett-Str. 40, 34132 Kassel, Germany, **Corresponding author **.

## Abstract

1

Endogenous circadian clocks control circadian rhythms in physiology and behavior. The predominant hypothesis of biological timing suggests that the responsible master clock for all endogenous circadian rhythms is constituted by an evolutionary conserved transcriptional-translational feedback loop (TTFL) clock consisting of positive feedforward and negative feedback elements. Unexpectedly, in contrast to the evolutionary derived insect *Drosophila*, RNAi-dependent knockdown of any of the negative feedback elements of the core TTFL clock in the basal Madeira cockroach *Rhyparobia maderae* does not delete circadian rhythms in locomotor activity. Shown here, neither RNAi-dependent triple knockdowns of all three negative feedback elements PERIOD, TIMELESS1, and CRYPTOCHROME2, nor single and double knockdown of the positive elements CLOCK and CYCLE did directly delete circadian locomotor rhythms as mRNA levels declined. Thus, our experimental data indicates the presence of compensatory elements, likely through posttranslational feedback loop (PTFL) controlled modifications. To explore alternative mechanisms, we constructed a computational model of a neuronal circadian pacemaker network using a network of coupled limit cycle oscillators, specifically planar switching affine systems (PSAS). The PSAS model comprises plasma membrane-associated PTFL clocks that are coupled to the TTFL nuclear clocks. Modeling results aligned with our experimental results. Therefore, both our experimental and modeling data support a systemic hypothesis of biological timing.

**Significance statement:** Based mostly upon genetic studies in derived taxa like *Drosophila* it is hypothesized that circadian timing of behavior is strictly controlled by specific circadian clock neurons in the brain, realized through a transcriptional-translational feedback loop (TTFL) clock. In contrast to this common hierarchical model that requires transcription, we provide first evidence in a basal taxon - the Madeira cockroach - for a systemic explanation of circadian timing of behavior that is based on coupled TTFL and posttranslational feedback loop (PTFL) clocks in adaptive neuronal networks.

## 4 Introduction

Endogenous physiological rhythms, such as circadian sleep-wake cycles or circannual activity rhythms, exhibit stable cycling on specific time scales. The ability to detect and predict external rhythms through synchronization of endogenous rhythms provides evolutionary advantages. In animals, physiological rhythms are orchestrated by dedicated neural circuits in the brain. These circuits produce endogenous oscillations on multiple physiologically relevant time scales and can be entrained to match periodic geophysical cues like light-dark cycles. In the mammalian brain, for example, the suprachiasmatic nucleus (SCN) is the circadian pacemaker center that drives sleep-wake cycles (Welsh et al., 2010). In insects, clusters of interconnected brain neurons were identified as circadian clock neurons (Brady et al., 2021; Helfrich-Förster, 1995; Reinhard et al., 2024; Reischig and Stengl, 2003a; Stengl and Homberg, 1994). In the Madeira cockroach, circadian clock neurons innervate a small neuropil in the optic lobes, the accessory medulla (AME), that serves as the circadian pacemaker center that controls sleep-wake cycles (Nishiitsutsuji-Uwo and Pittendrigh, 1968; Reischig and Stengl, 2003a; Stengl et al., 2015; Stengl and Homberg, 1994).

### 4.1 Circadian pacemaker centers comprise a morning and an evening oscillator circuit

The circadian pacemaker center consists of ensembles of coupled neuropeptidergic clock neurons that control different physiological functions such as sleep or wakefulness. However, endogenous circadian rhythms do not start at the network level, they originate at the level of single clock cells. Individual AME and SCN clock neurons generate endogenous circadian action potential rhythms (Jackson et al., 2004; Schneider and Stengl, 2007; Webb et al., 2009; Welsh et al., 1995).At defined zeitgeber times, action potential activity peaks, which rhythmically releases neuropeptides like PDF or VIP to recruit other clock neurons into synchronized ensembles that time physiological contexts, such as sleep-wake cycles (Aton et al., 2005; Helfrich-Förster, 1995; Helfrich-Förster et al., 2000; Patton et al., 2020; Schneider and Stengl, 2005; Stengl and Arendt, 2016). At least two synchronized oscillator ensembles of clock neurons exists in the circadian pacemaker center of mammals and insects that track the length of the daily light phase (photoperiod) (Helfrich-Förster, 2009; Jagota et al., 2000; Pittendrigh and Daan, 1976): The morning oscillator (MO) tracks dawn and the evening oscillator (EO) tracks dusk. In the crepuscular *Drosophila* its bimodal activity pattern with peaks at dusk and dawn (Helfrich-Förster, 2000) is hypothesized to originate from antiphase coupling of the MO and EO clocks (Grima et al., 2004; Helfrich-Förster, 2001; Pittendrigh and Daan, 1976; Stoleru et al., 2004).

### 4.2 Circadian clock neurons express conserved core clock genes

Like all clock neurons, MO and EO clock neurons are characterized by the circadian expression of core clock genes that constitute a transcriptional/translational feedback loop (TTFL) (Hastings et al., 2019). In insects, the TTFL clock is best studied in the fruit fly *Drosophila melanogaster*. There, the transcription factors CLOCK (CLK) and CYCLE (CYC; homologue of BMAL1 in vertebrates) form the positive feedforward elements. They heteromerize and activate the transcription of the negative feedback genes *period* (*per*) and *timeless1* (*tim1*). The respective proteins PER and TIM1 close the loop via delayed inhibition of their own transcription, thus generating circadian rhythms in mRNA- and clock protein levels (Hardin, 2000). There is growing evidence that the circadian clock of the nocturnal Madeira cockroach also comprises a MO and EO that oscillate in antiphase, despite the absence of a bimodal activity pattern. While the TTFL clock of the Madeira cockroach shares elements with *Drosophila*, it employs CRYPTOCHOME 2 (CRY2) as additional negative feedback element, similar to the mammalian TTFL clock (Tomioka and Matsumoto, 2010; Werckenthin et al., 2012). Furthermore, in the cockroach, each clock circuit appears to be based on a network of MO or EO clock neurons with differential gene expression: Neurons of the MO project to the ipsilateral AME and are hypothesized to express *per*, while neurons of the EO project to the ipsi-and contralateral AME and are hypothesized to express *per*, *cry2*, and *tim1* (Gestrich et al., 2018; Werckenthin et al., 2020). The hypothesis of two clock circuits per AME is further supported by the bimodal peaks of cAMP levels in the optic lobes. cAMP peaks at dusk and dawn suggest that there are two circadian clock ensembles per AME that are synchronized in antiphase by PDF increasing cAMP levels in respective clock neurons (Gestrich et al., 2018; Schendzielorz et al., 2014; Wei et al., 2014).

### 4.3 Different types of clocks can control circadian rhythms in physiology and behavior

In animal chronobiology it is generally assumed that, in a hierarchical model, the circadian TTFL clock of master clock neurons in the circadian pacemaker center of the brain dictates circadian rhythms in behavior as direct TTFL clock output (Hastings et al., 2019; Honma, 2018; Patton and Hastings, 2023). Accordingly, in *Drosophila,* deletion of any of the circadian clock genes of either the negative or the positive elements of the TTFL clock in circadian clock neurons obliterated circadian locomotor activity rhythms (Allada et al., 2003; Konopka and Benzer, 1971; Rutila et al., 1998; Sehgal et al., 1994). However, knockdown of individual negative elements comprising the TTFL clock of circadian clock neurons in the Madeira cockroach did not cancel circadian locomotor behavior but only changed its period (Werckenthin et al., 2020). Thus, there may be additional periodic posttranslational mechanisms comprising delayed negative feedback elements (oscillators and clocks) besides the TTFL clock in the cockroach, similar to those in cyanobacteria that rely on a posttranslational feedback loop (PTFL) clock of autonomous protein phosphorylations to produce circadian rhythms (Nakajima et al., 2005; Tomita et al., 2005). Another TTFL-independent clock is based on circadian redox cycle, such as human red blood cells, which lack a nucleus and DNA (O’Neill and Reddy, 2011).

### 4.4 Our novel hypothesis of systemic timing in the Madeira cockroach

Therefore, for timing in the cockroach, we propose an alternative, systems-based model of biological timing in contrast to the hierarchical model. We predict many flexibly coupled TTFL and PTFL oscillators / clocks in each clock cell, constituting an adaptive clock system. One feature of such a heterarchical system of distributed, adaptive pacemaker networks of clock neurons would be to flexibly synchronize with various environmental rhythms (Stengl and Schneider, 2024; Stengl and Schröder, 2021).

Here, we used a combination of experimental and computational modeling studies to study the limits of the TTFL clock in the long-lived large Madeira cockroach *Rhyparobia maderae*, an established cellular model of chronobiology from a more basal taxon than *Drosophila* (Nishiitsutsuji-Uwo and Pittendrigh, 1968; Petri and Stengl, 1999). To expand on previous studies (Werckenthin et al., 2020, 2012), we first knocked down all negative feedback elements of the cockroach TTFL clock simultaneously, as well as both or single positive elements. Subsequently, we compared the time course of the decrease of the respective mRNA levels of *Rm’clk* gene expression with the development of arrhythmic locomotor activity in running wheel assays. If compensatory mechanisms for the reduction of clock proteins exist, changes in circadian locomotor activity would not correlate with mRNA levels. Last, we corroborated our systemic hypothesis with a mathematical model to allow predictions of manipulations that delete circadian locomotor rhythms in the Madeira cockroach.

We propose a novel computational modeling scheme for the circadian clock network in the Madeira cockroach to simplify and enhance analysis and interpretation of the biological experimental results. Single Goodwin or van-der-Pol oscillators are widely used in biology to describe a wide variety of oscillations (Escalante-Martínez et al., 2018; Goldbeter, 2002; Gonze and Ruoff, 2021) including the human circadian clock (Tayong et al., 2022). The ordinary differential equations underlying these oscillations are continuous but essentially (strongly) nonlinear. This makes the analysis of stability, robustness, or the sensitivity of periods or phase shifts to model parameters very challenging, leaving often only numeric simulation as a means for model evaluation, but preventing analytic solutions. Here, we selected the class of planar switching affine systems (PSAS) to represent a single periodic phenomenon, such as the production and degradation of a single core clock protein. These single oscillators were interacting in a network structure to represent relevant effects of the biological clock. Here, the states of the PSASs represent cellular substances, such as mRNA levels and cAMP concentrations. The network structure models the mechanistic interactions within the clock neuron – between its nucleus and plasma membrane – as well as the interactions between clock neurons. PSAS have recently been introduced in Hanke and Stursberg (2023) as oscillator class consisting of affine subsystems assigned to polyhedral regions of the partitioned state-space. For suitable parameterization, the oscillations are obtained as alternating execution of affine dynamics with intermittent switching, which allows for the derivation of explicit analytical solutions. In addition, the adaptation of this model class to experimental data is relatively straightforward (Hanke et al., 2024). Since PSAS oscillators retain linear structure within each mode, linear system theory becomes applicable, thus enabling a tractable analysis of robustness, of the effects of external signals, or synchronization. The coupling of several PSAS oscillators forms networked structures of interacting and synchronizing rhythms, which serve to represent and explain the principles of the underlying biological clocks.

## 5 Methods

### 5.1 Animals

*Rhyparobia maderae* cockroaches were bred and raised in mass colonies at the University of Kassel under 12 hours light/12 hours dark (LD 12:12) conditions, 25°C and 50% relative humidity. Housing consisted of large translucent plastic bins (60×40×30 cm), filled with litter and cardboard egg cartons. Bins were covered with metal mesh and a 5 cm Vaseline rim to prevent animals from escaping. Cockroaches were fed with dried dog food, organic potatoes and water *ad libitum*. We used only adult male animals in this study because circadian locomotor activity is more variable in females and masked by the reproductive cycle (Leuthold, 1966; Tsai and Lee, 2000; Wiedenmann and Martin, 1980). Furthermore, this ensures comparability and continuity with previous studies.

For age-controlled experiments, freshly molted males were separated from the colony and kept in smaller holding boxes. Depending on the number of animals per cohort, groups of 10 - 20 individuals were housed in transparent plastic boxes (27×18×17 cm) and groups of 20 - 40 individuals in larger boxes (31×18×12 cm). Animals designated for LD were kept under the same light conditions as their source mass colony. Animals for experiments in constant darkness (DD) were transferred to constant darkness immediately after separation.

### 5.2 Running wheel assay

We observed the free-running locomotor activity of individual cockroaches in custom-built running wheels in DD and constant light (LL) conditions, 25°C and 50% relative humidity. In the running wheels, cockroaches were isolated from each other, but 16-20 wheels were housed together in a light-tight recording box. Even though this enabled the exchange of chemosensory cues with box mates to mimic colony housing conditions, which could potentially act as Zeitgeber, we did not observe synchronization of locomotor activity across individuals within a recording box (Supplementary Figure S 1). Animals had access to food and water *ad libitum*.

The wheels were constructed from transparent plastic with an outer diameter of 7 cm and an inner ring of 3.5 cm diameter, providing a running channel of approximately 3.5 cm width for the animals. The outer ring featured evenly spaced perforations of approximately 4 mm diameter, spaced ∼1.8 mm apart, allowing frass and debris to fall through without impeding locomotion. The plastic wall itself had a thickness of ∼1mm. Two magnets were mounted on the outer rim at 180° intervals, allowing each half-rotation to be detected by a Hall sensor. Activity was recorded via two custom Arduino-based logging systems. In the standard configuration, events were sampled and stored directly in 1 min bins onto an SD card. In the timestamp configuration, each Hall sensor trigger was recorded with a timestamp at 1-second resolution; data were collected from both Arduino on a Raspberry Pi and extracted as plain text files, then binned into 1 min intervals for analysis.

Data were visualized as double-plotted actograms with a bin size of 10 min. Actogram bar height was scaled logarithmically to the number of running wheel turns per bin, capped at the median activity of non-zero bins, such that bins at or above the median are displayed at full height.

### 5.3 RNA interference (RNAi) and knockdown efficiency

Only animals that showed rhythmic behavioral activity (see Data analysis below) for at least 7 days in DD were used for RNAi experiments. If animals were recorded for less than 10 days, missing data was filled in with zeros to meet the 10-day window criterion (see below). We used double-stranded RNA (dsRNA) for the positive elements *Rm’clk* and *Rm’cyc* and the negative elements *Rm’per*, *Rm’tim1*, *Rm’cry2* of the molecular TTFL clockwork. dsRNA was prepared and injected as described in (Werckenthin et al., 2020). Primer sequences are listed in Table 1. In brief, 6 μg dsRNA per target gene was dissolved in 6x concentrated, sterile filtered saline (concentration in mM: 768 NaCl, 16.2 KCl, 12 CaCl_2_, 7.2 NaHCO_3_, pH = 7.5), and filled with nuclease-free water to 4 μl, resulting in a final saline concentration of 1x. Animals were injected with either 6 μg dsRNA of *gfp* as a control, because GFP is not present in the cockroach, or the targets *Rm’per*, *Rm’tim1*, *Rm’cry2*, *Rm’clk* and *Rm’cyc.* We used 6 μg of each target dsRNA, thus total dsRNA was 6 µg for single knockdowns, 12 µg for double knockdowns (DKD), and 18 µg for triple knockdowns (TKD).

**Table 1:**
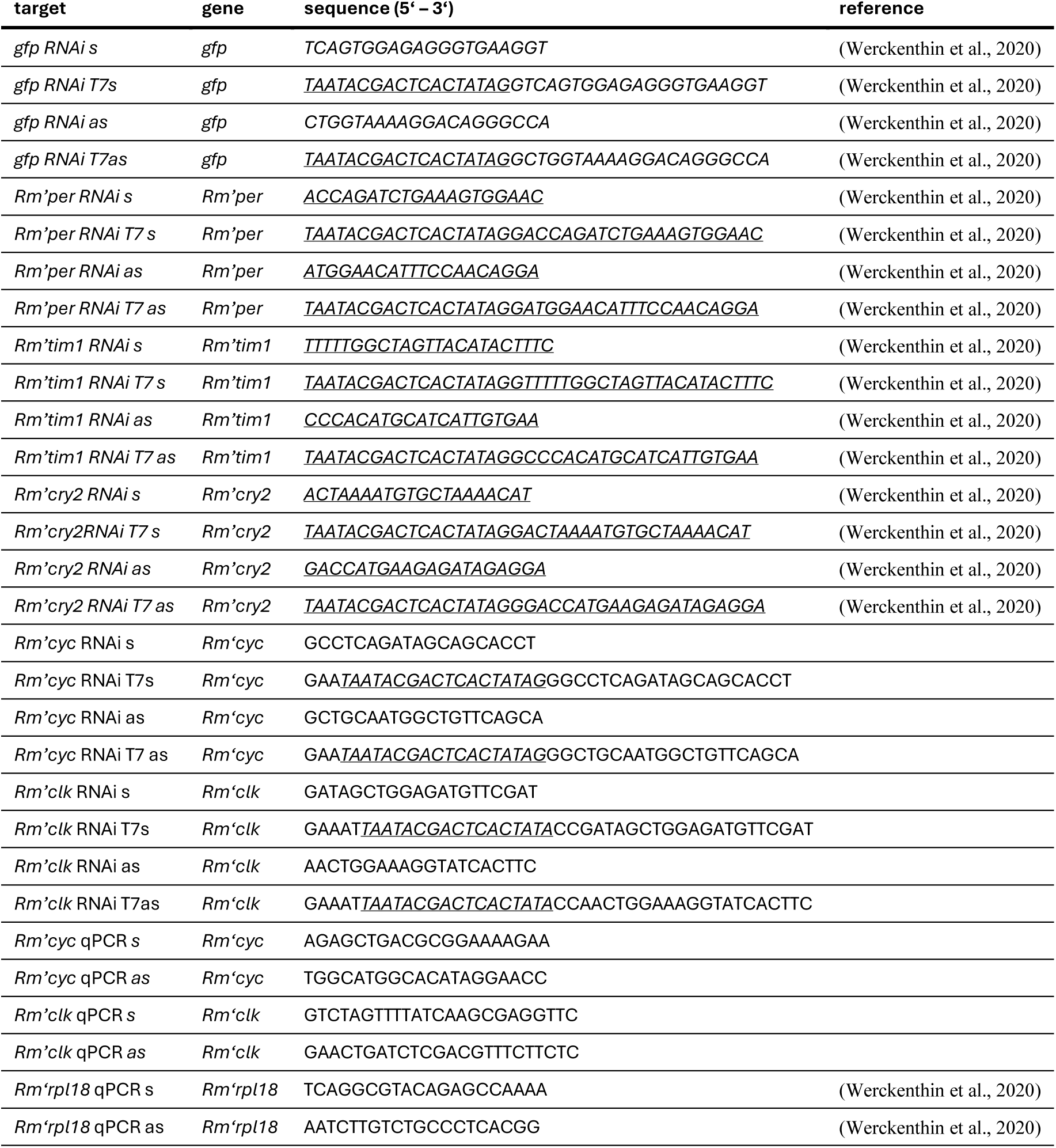
Primer sequences used for dsRNA generation and qPCR targets. T7 sequence is underlined.

Animals were cold-anesthetized by immersion in ice-cold water for 30 s and injected ventrally at the thorax joint between the pro- and mesothoracic leg pairs (Werckenthin et al., 2020). Immediately after injection, locomotor activity was monitored in running wheels for two subsequent months. To test the efficiency of RNAi knockdowns, dsRNA-treated animals from DD conditions were used for RNA extraction with subsequent qPCR (see below; primers in Table 1) at projected ZT4 in a time series after 2, 3, 4, 5, 6, 7, 14, 21, and 60 days, respectively (*Rm’clk* knockdown), 28 days (*Rm’cyc* knockdown), or 55 days (TKDs). To control for potential age-related decrease in mRNA levels, we transferred age-controlled males 2 weeks after their final molt into DD. After 2 days in DD, we measured relative mRNA levels with qPCR as control (t_0_), and 7, 14, and 23 days after t_0_, corresponding to the same time window in which the time series showed a significant decrease of *Rm’clk* mRNA levels in the respective knockdown animals.

### 5.4 Daily and circadian mRNA time series

To investigate daily and circadian oscillations of *Rm’clk* and *Rm’cyc* mRNA levels, we collected samples of untreated animals 3-4 weeks after the final adult molt. Samples were collected in 30 min windows, centered on the respective ZT timepoint (ZT 0, 4, 8, 12, 16 and 20). Animals were anesthetized by immersion in ice-cold water for 30 s immediately before dissection. To measure daily mRNA levels, animals were collected from the LD colony immediately before dissection. To measure circadian oscillations with minimal dispersion of the free-running rhythms, animals were first transferred to constant darkness at ZT 12 and collected 24-48 hours later: We collected samples at ZT 12, 16, and 20 during the first circadian cycle, and ZT 0, 4, and 8 during the second circadian cycle (Figure 1). Animals from ZT 0 were dissected under dim red light, and animals from ZT 12 were dissected in normal light, the other timepoints were dissected in their respective light condition, either red dim light or normal light.

**Figure 1:**
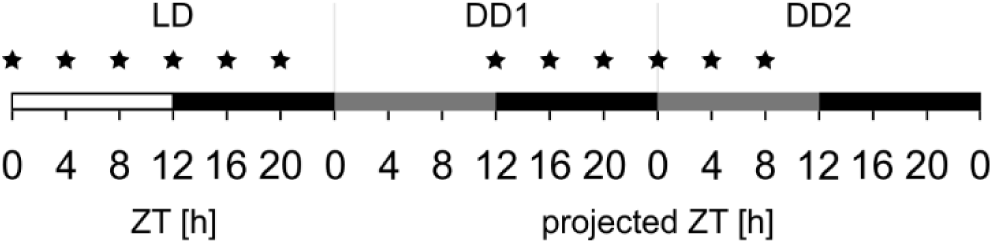
Sampling scheme for cycling of *Rm’clk* and *Rm’cyc* in LD and DD. Stars in LD phase indicate sampling points for LD experiment; stars in DD1 (first day of DD) and DD2 (second day of DD) indicate sampling points for DD experiments.

For each timepoint, three brains were extracted, pooled, and snap-frozen in liquid nitrogen. For each ZT, we obtained 10 pools of 3 brains in LD and 6 pools of 3 brains in DD. Each pool was measured in triplicate in the quantitative real-time polymerase chain reaction (qPCR) analysis (see below).

### 5.5 Quantitative real-time PCR

RNA was extracted from whole brains using NucleoSpin® RNA (Machery-Nagel) according to the manufacturer’s protocol. qPCR was conducted using 4X CAPITAL™ 1 Step qRT-PCR Probe Master Mix (biotechrabbit) according to the manufacturer’s protocol, together with the Eppendorf Mastercycler® RealPlex^2^ (Eppendorf) in 96-well PCR plates (white; Biozym Scientific). Single amplicons were confirmed with melting-curve analysis and sanger sequencing (Microsynth Seqlab). As reference gene, we used ribosomal protein L18 (*Rm’rpl18*: MT524704), which is expressed at stable levels in many insects across tissues, developmental stages, and sexes (Gong et al., 2024). However, during data analysis we detected significant oscillations in *Rm’rpl18* Ct values across ZT timepoints (one-way ANOVA: LD, F(5, 54) = 7.11, p < 0.001; DD, F(5, 30) = 5.90, p < 0.001), which has not been reported before. We therefore opted for grand-mean normalization, in which expression values are normalized to the mean Ct of all timepoints per gene per condition, rather than conventional ΔΔCt normalization.

### 5.6 Data analysis

#### Statistics

Analysis and data visualization of the behavioral experiments and mRNA time series were performed using R 4.3.0 and RStudio (2023.03.1). All statistical analyses were performed in R with emmeans package (Lenth and Piaskowski, 2025). When data was normally distributed (Shapiro-Wilk test p > 0.05), parametric tests were applied; otherwise, non-parametric equivalents were used as indicated. Statistical significance was assessed at α = 0.05 and is indicated as * p < 0.05, ** p < 0.01, *** p < 0.001.

#### Locomotor activity

Based on the observed locomotor patterns, behavioral data was split into three phases with dsRNA injection on day 0: 1) pre-knockdown phase (pre-phase) day -10 to -1 before injection; 2) transition phase (trans-phase) day 0 to 13 after the injection to account for transient treatment effects; 3) post-knockdown phase (post-phase) day 14 – end of recording after the injection, when behavioral activity had reached a new steady state. Rhythmicity of the locomotor activity was evaluated after Reischig and Stengl (2003b) for each pre-, trans-, and post-phase for each animal. In brief, the peak of a χ²-periodogram, normalized to the Sokolove-Significance Line (SSL), must exceed that line by 20 % and have a width of ≥ 0.7 (bin size: 30 min, window size: 10 days). The pre-phase consisted of one window. For the trans- and post-phase, the 10-day window was advanced by 1 day until the beginning of the next phase or end of the recording, respectively. Each window was evaluated for rhythmicity, and the percentage of rhythmic windows was calculated for the trans- and post-phases for each animal. Rhythmicity percentages were compared between knockdown groups using a Wilcoxon rank-sum test since normality assumptions were not met (Shapiro-Wilk test). Animals were classified as arrhythmic if they showed no rhythmic windows within the post-phase between day 14-50, which is a more conservative criterion than previously used (Werckenthin et al., 2020). For rhythmic animals, period length was estimated from 30-minute binned data using a 5-day sliding-window cosinor fit (step size 0.5 days; period search range 20–28 h in 0.1 h increments) including zero-amplitude test. The median period was taken for each animal across all windows in the pre- and post-phase, respectively, that yielded significant cosinor fits.

TKD animals were classified as period-lengthened or period-shortened if the difference between pre-and post-phases for the individual exceeded ±0.5 h; otherwise, they were classified as unchanged. For single and double knockdowns, periods were compared across pre-, trans-, and post phase with one-way repeated measures ANOVA and Tukey’s post-hoc test, unless indicated otherwise.

Recovery of rhythmicity during the last 10 days of recording in *Rm’clk*-KD and DKD animals was assessed using the 5-day sliding-window cosinor fit described above. An animal was classified as rhythmic if the fit was significant and R^2^ > 0.159 within at least 6 consecutive windows (spanning 3 days) and a gap tolerance of up to 2 non-passing windows (1 day). The R² threshold was defined as the 25^th^ percentile of R² values from all significant pre-phase windows across all animals in this study (n = 83), providing an empirical lower bound for a genuine circadian cosinor fit.

#### mRNA time series

For time series expression of mRNA levels in wild-type animals, all data points were retained as natural biological variation. *Rm’clk* and *Rm’cyc* mRNA levels were normalized to the grand mean of the reference gene *Rm’rpl18* across all ZTs. Circadian and ultradian rhythmicity was tested with cosinor regression by fitting separate sinusoidal models with periods of 24 h and 12 h. Differences in relative mRNA expression across timepoints were assessed with one-way ANOVA and Tukey’s post-hoc test for multiple comparisons.

Knockdown efficiency was assessed by measuring relative mRNA levels of *Rm’clk* at 2, 3, 4, 5, 6, 7, 14, 21, and 60 days post-injection. Differences across timepoints were tested using Kruskal-Wallis test with Dunn’s post-hoc test and Holm correction. Knockdown efficiency for the other core clock genes was assessed by comparing expression levels of the respective mRNA to control animals injected with dsRNA of *gfp* at the same time point, or by comparing mRNA of injected animals to untreated controls with parametric or non-parametric tests as indicated.

### 5.7 Computational model of the cockroach circadian clock network

To construct a computational model of the cockroach circadian clock under DD conditions, we first simplified the network to two distinct oscillators (Pittendrigh and Bruce, 1957), one leading morning (MO; period < 24 h) and one lagging evening (EO; period > 24 h) oscillator, to characterize the rhythmic alternation between activity and resting, reminiscent of the morning and evening oscillators in *Drosophila* (Stoleru et al., 2004) (Figure 2A). Here, all biological neurons belonging to the MO are represented by the computational MO cell, and all neurons of the EO are represented by the EO cell. While the terms ‘lead’ and ‘lag’ are commonly used in control engineering by a well-established concept where they refer to the effects of lead and lag compensators, i.e., phase-advancing and phase-delaying actions, respectively (e.g., (Franklin et al., 2019)), we will use the terms MO and EO that are better established in chronobiology throughout this manuscript. When coupled, the MO and EO cells shall synchronize in period but have a constant phase shift.

**Figure 2:**
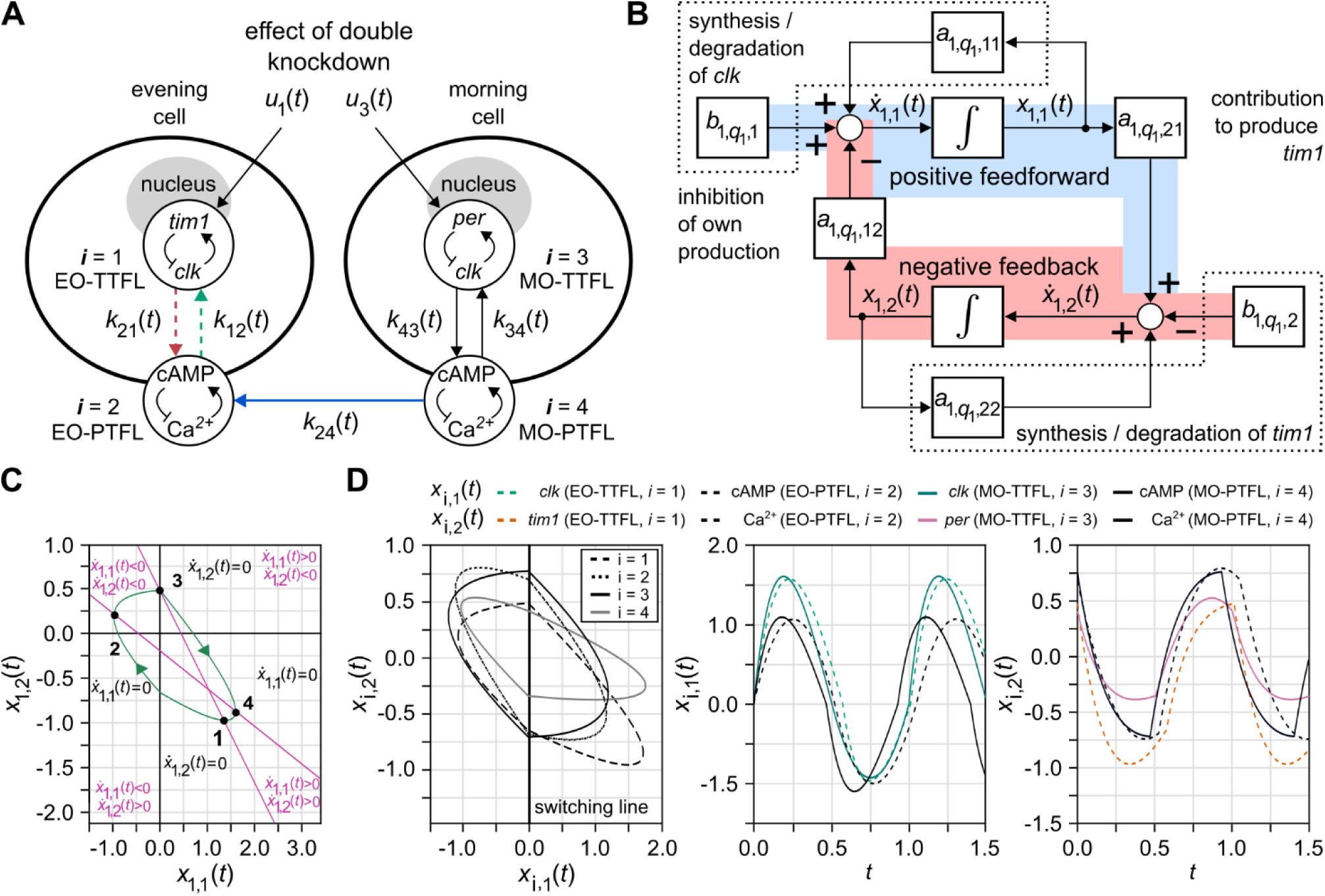
The PSAS model of the cockroach clock network that determines the behavior of the animal in constant darkness (DD). **(A)** The network models the DD condition where cockroaches express a free-running locomotor activity rhythm with a period of approximately 22.8 h, in which the MO (cycle period <24 h) dominates the EO (cycle period >24 h). MO and EO are each represented by one cell. Each cell comprises one TTFL (*i* = 1 and 3) and one PTFL (*i* = 2 and 4) oscillator with bidirectional coupling. Both cells are coupled via their PTFLs. Since the MO cell dominates in the condition modeled here, the coupling gain (*k(t)*) from the EO-PTFL to MO-PTFL was set to zero and is not shown in this schematic. The EO-TTFL models oscillations of *clk* and *tim1* levels, the MO-TTFL of *clk* and *per* levels (Werckenthin et al., 2020). Both PTFLs model the oscillations of Ca^2+^ and cAMP levels. They are linked to but not forced by the TTFLs. The input *u*(*t*) represents the knockdown of the *Rm’clk* gene and is therefore acting on both TTFLs. Gain changes for the colored arrows are shown in Figure 7A. **(B)** Block diagram of the biological feedforward and feedback mechanisms within the EO-TTFL (*i* = 1): Blocks represent the amplification of an incoming signal with a model parameter of the PSAS, or the integration of the signal over time; circles model the addition or subtraction of signals. **(C)** Clockwise oscillation of the limit cycle in the EO-TTFL. Synthesis of *clk* and *tim1* occurs when *ẋ*_1,1_(*t*) > 0 and *ẋ*_1,2_(*t*) > 0, while degradation of *clk* and *tim1* occurs when *ẋ*_1,1_(*t*) < 0 and *ẋ*_1,2_(*t*) < 0. The change of sign on the limit cycle is illustrated by the magenta lines through points 2 and 4 and points 1 and 3, respectively. **(D)** Local limit cycles of all four oscillators in phase space (left panel) and time domain (middle and right panel). The solutions *x_i_*(*t*) switch when they reach the switching line, which contributes to the evolution of the limit cycles.

Within each model cell, a coupled pair of TTFL and PTFL oscillators drive the cell oscillation. To further reduce complexity, the MO-TTFL comprises the interacting clock genes *clk* and *per*, whereas the EO-TTFL comprises oscillations of interacting *clk* and *tim1* (Werckenthin et al., 2020). Since *cyc* oscillates similar to *clk* it was neglected. While our understanding of the PTFL is still sparse, experiments have demonstrated antiphase oscillations of the important 2^nd^ messenger molecules cAMP and Ca^2+^, with peak cAMP levels at dusk and dawn and peak Ca^2+^ levels during the day (Rojas et al., 2019; Schendzielorz et al., 2014). Therefore, in the model the two PTFL oscillators are simplified to the interaction between cAMP and Ca^2+^. For the MO-PTFL, a period of 23 h is assumed and a lengthened period of 25 h for the EO-PTFL.

Within the MO and EO cells, the respective TTFL and PTFL are bidirectionally coupled. Additionally, MO and EO cells are coupled to each other via their respective PTFLs. Since the free-running circadian period of cockroaches shortens in DD conditions, we assume that the MO oscillator dominates the behavior by inhibiting the EO oscillator via GABA release (Werckenthin et al., 2020). Thus, we omitted the connection from EO-PTFL to MO-PTFL in this model.

The clock networks is formulated by coupled planar switching affine system (PSAS) oscillators (Hanke et al., 2024), where each of the four single oscillators with index *i* ({1, …, 4}: EO-TTFL, EO-PTFL, MO-TTFL, MO-PTFL) is modeled by

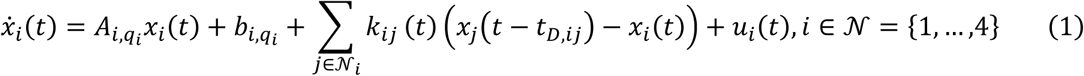

with *x_i_*(*t*) ∈ ℝ^2^ defining the state, *q_i_* ∈ {1,2} the mode, *A_i_*_,*q*_*_i_* ∈ ℝ^2×2^ the system matrices, and *b_i_*_,*q*_*_i_* ∈ ℝ^2^ the affine terms. The coupling term is denoted by

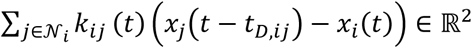

with a dead time (time delay) *t_D_*_,*ij*_ ∈ ℝ_≥0_, *N_i_* ⊂ *N* determines the set of indices of neighboring oscillators and *u_i_*(*t*): ℝ ↦ ℝ^2^introduces an additional input for *i* ∈ {1,3} to model knockdown. The gain *k_ij_*(*t*) associates with the coupling of the oscillators with indices *i* and *j* chosen to be time-varying to suitably model changes in reaction to knockdown experiments. The dynamics assigned to *q_i_* = 1 is active for

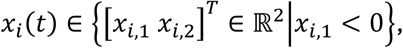

while the dynamics for *q_i_* = 2 is active in the half-plane

*x_i_*(*t*) ∈ {[*x_i_*_,1_ *x_i_*_,2_]*^T^* ∈ ℝ^2^|*x_i_*_,1_ > 0}. The uncoupled local oscillators converge to unique and stable limit cycles if the parameters *A_i_*_,*q*_*_i_* and *b_i_*_,*q*_*_i_* satisfy specific construction rules (see (Hanke et al., 2024)). The most important rule is that the steady states *x̅_qi_* ∈ ℝ^2^ lie in the opposite half-plane, so that the states *x_i_*(*t*) never reach them. *x_i_*_,1_ = 0 determines the switching line, which causes switching between the two dynamics, as illustrated in Figure 2D. In contrast to other oscillators, such as the van-der-Pol (Forger et al., 1999) or Goodwin oscillators (Goodwin, 1965), which are restricted in the form of their limit cycles, the limit cycle of a PSAS can be very flexibly adjusted to measured data. The states for the oscillators are defined as follows:

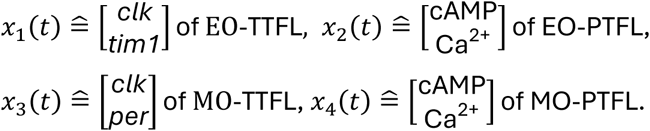

A heterarchical network can be represented by the previously defined network, in which every oscillator consists of a PSAS and choosing the coupling gains from TTFL to PTFL and from PTFL to TTFL to be similar. In the heterarchical network, both TTFLs and PTFLs constitute endogenous clocks.

Similar to Goodwin oscillators (Goodwin, 1965), the parameters of a PSAS oscillator can be interpreted as synthesis, production or inhibition of a biological quantity, such as mRNA. For the EO-TTFL oscillator, comprising *clk* and *tim1*, these interpretations and the resulting feedforward and feedback pathways are illustrated in Figure 2B. Here, *x*_1,1_(*t*) = *clk* resides in the feedforward pathway, while *x*_1,2_(*t*) = *tim*1 belongs to the feedback pathway. The combination of the terms *a*_1,*q*1,11_ and *b*_1,*q*1,1_ results in either degradation or transcription/translation (synthesis) of *clk*, depending on which half-plane (*q*_1_ = 1 or *q*_1_ = 2) the state *x*_1_(*t*) currently evolves. The sign of the affine term and *a*_1,*q*1,11_(*t*)*x*_1,1_(*t*) changes. A similar mechanism applies to the synthesis of *tim1*, where the expression *a*_1,*q*1,22_*x*_1,2_(*t*) − *b*_1,*q*1,2_ determines whether synthesis or degradation occurs. To adapt the PSAS parameters to the data, the data values are normalized such that they oscillate around zero within a range of −1 to 1.

The degradation and synthesis of *clk* and *tim1* modeled by the PSAS is further illustrated in Figure 2C: Consider the segment of the limit cycle from point 1 to 2 for which *ẋ*_1,1_(*t*) < 0 and *ẋ*_1,2_(*t*) > 0: *clk* is degraded, while *tim1* is synthesized. This reflects the inhibitory effect of *tim1* on *clk*. Beyond point 2 in direction of the limit cycle, synthesis of *clk* resumes with *ẋ*_1,1_(*t*) > 0, though at a reduced rate, while synthesis of *tim1* slows down, decreasing with *ẋ*_1,2_(*t*) but remaining positive up to point 3. Between the points 3 and 4, *clk* accumulates (*ẋ*_1,1_(*t*) > 0), while *tim1* undergoes degradation (*ẋ*_1,2_(*t*) < 0) and is thereby disinhibiting *clk*. Beyond point 4, *clk* reaches high levels, driving the synthesis of *tim1* (*ẋ*_1,2_(*t*) > 0).

To model the interconnection of different oscillators, an often-used approach for synchronizing oscillators is diffusive coupling, in which the sum of the differences between neighboring oscillators is weighted by a coupling gain, as defined in Eq. (1). As the coupling gain increases, the oscillations gradually synchronize. Since the PTFL and TTFL oscillators jointly govern the cellular rhythms in MO and EO cells, it is straightforward to apply this coupling scheme for the pairs (*i*, *j*) ∈ {(1,2), (2,1), (3,4), (4,3)} without a dead time *t_D_*_,*ij*_. For (*i*, *j*) = (2,4), a dead time is used to bring these oscillators in antiphase. This is realized by coupling mechanism between the MO-PTFL and EO-PTFL. Therefore, *t_D_*_,24_ is set to half of the period of the MO-PTFL, i.e., *t_D_*_,24_ = 0.9325/2 days. The selection of coupling gains is closely tied to the analysis of interactions between oscillators, leading to the question of how large the impact of the coupling gains is on the behavior of single oscillators. Given that molecular and cellular interactions are not well established, the determination of coupling gains aids in identifying connections that are essential for a functional biological clock. Therefore, the coupling gains were adjusted in a stepwise manner, beginning from zero, while accounting for mutual influences, namely different combinations of gains, and aiming toward a heterarchical network. The most suitable combination of gains is presented in the Results.

To capture the effect of knocking down *clk*, the input variable *u_i_*(*t*) must be defined. To this end, let *t_k_* ∈ ℝ_>0_ denote the time at which *clk* knockdown is initiated. The following biological observations (see Results) are included in the model for selecting *u_i_*(*t*) for *i* ∈ {1,3}:

- For *t* ∈ [0, *t_k_*], the MO and EO oscillators are in antiphase, reflecting the normal rhythmic behavior.
- For *t* ∈]*t_k_*, *t_k_* + 7], the mRNA levels drop nearly to zero, leading to vanishing oscillations of the TTFLs, i.e., amplitudes decrease over time, and the period of the TTFLs (together with the behavioral activity patterns) gradually increases, and the interaction between the two PTFL oscillators gradually diminishes in consequence.
- For *t* ∈]*t_k_* + 7, *t_k_* + 14], the coupling from the TTFLs to the PTFLs reduces upon *clk* knockdown, as *clk* regulates the strength of this connection. Thus, the coupling gains from the TTFLs to the PTFLs remain unchanged, since the strength of these couplings is scaled relative to the magnitude of random fluctuations around zero and the difference between neighboring states. Under these conditions, the interaction between the PTFL oscillators is assumed to diminish toward a value close to zero at *t* = *t_k_* + 14.
- For *t* ∈]*t_k_* + 14, *t_k_* + 40], the cockroaches exhibit arrhythmic behavior; the oscillators lose synchronization and no coherent rhythmic output is observed.
- For *t* ∈]*t_k_* + 40, ∞[, rhythmic behavior reemerges without recovery of mRNA levels, indicating that the oscillators (partially) resynchronize, i.e. the PTFL oscillators synchronize in antiphase, whereas the TTFL oscillators exhibit only weak rhythmic activity. This suggests that *clk* and *cyc* knockdown has caused permanent dysfunction of the TTFLs. Furthermore, the knockdown signal remains active, maintaining the disrupted state. The underlying biological mechanism has not yet been identified.

To capture these effects in modeling, the steady states *x̅_qi_* are shifted towards the switching line upon knockdown. This shift lengthens the period of the PSAS oscillators, as trajectories move more slowly close to steady states. Simultaneously, the amplitudes decrease because the trajectories travel shorter distances as the steady states approach the switching line. To represent spontaneous, low-amplitude fluctuations in protein levels, a white noise signal *σ*(*t*) ∈ ℝ, ∀*t* > *t_k_* + 7 is introduced, reflecting random *clk* expression. *σ*(*t*) is applied uniformly to both components of *ẋ*_1_(*t*) and *ẋ*_3_(*t*), as the entire TTFL is influenced by the mutual dependence of *clk* and *tim1*, respectively *per*. In this regime, the TTFL oscillators exhibit random fluctuation around the origin. At *t* = *t_k_* + 7, the input switches to *σ*(*t*)1_2_ − *b_i_*_,*q*_*_i_*, and a random oscillation around zero is obtained by reducing the uncoupled dynamics to *ẋ_i_*(*t*) = *A_i_*_,*q*_*_i_x_i_*(*t*) + *σ*(*t*), for *t* ∈]*t_k_* + 7, ∞[. Thus, the knockdown effect is modeled for *i* ∈ {1,3} according to:

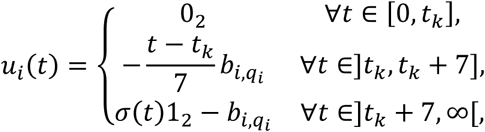

where 1_2_ (0_2_) denotes a two-dimensional vector of ones (zeros).

In addition, we created a hierarchical model with dominant TTFLs in a similar manner with two key differences: 1) the PTFLs do not constitute endogenous clocks but act as followers whose oscillations are driven by the TTFLs; 2) coupling from the dominant TTFL to the PTFL is stronger than the coupling from the PTFL to the TTFL. The uncoupled dynamics of the PTFLs (*i* ∈ {2,4}) are now modeled by the linear, transient system

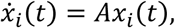

where *A* is assumed to be a Hurwitz matrix, i.e., all eigenvalues of *A* have strictly negative real part. Thus, the state of the uncoupled PTFL will converge to 0_2_ for *t* → ∞. The coupling gains of the hierarchical network are chosen in a similar manner as for the heterarchical network.

## 6 Results

Knocking down individual key elements of the negative limb (*Rm’per*, *Rm’tim1*, or *Rm’cry2*) of the molecular clock in the Madeira cockroach does not abolish circadian locomotor activity (Werckenthin et al., 2020). This hints at redundancies in the clockwork. One possibility is that the other, unaffected negative elements could compensate for the one knocked down negative element. Alternatively, in addition to the TTFL clock, posttranslational feedback-controlled mechanisms (PTFL clocks) could ensure functional, circadian electric output of clock neurons as long as sufficient supply of proteins like ion channels and postsynaptic receptors is available. In this study, we explore these possibilities in the Madeira cockroach by expanding on our previous work. First, we simultaneously knocked down all three elements of the negative limb of the cockroach TTFL clock. Second, after ensuring daily and circadian cycling of the positive elements *Rm’clk* and *Rm’cyc*, we knocked them down individually or simultaneously. Finally, because the time course of declining mRNA levels did not correlate with the observed circadian locomotor output, we applied a computational model of coupled TTFL and PTFL oscillators as planar switching affine systems (PSAS) to investigate whether oscillatory output of clock neurons that represents circadian locomotor activity is still possible with PTFLs only, after eliminating the TTFL clock, and used the model to predict the behavior of cockroaches in LL before and after knockdown.

### 6.1 Triple knockdown of negative TTFL elements does not disrupt rhythmic activity

Since single knockdowns of *Rm’per*, *Rm’tim1* and *Rm’cry2* do not result in loss of circadian locomotor rhythmicity in the Madeira cockroach (Werckenthin et al., 2020), we simultaneously targeted all three known negative elements in a triple-knockdown (TKD) (Figure 3). After recording 10 days of running wheel activity in DD as control (“pre”-phase, day -10 – -1), we injected animals that expressed robust circadian rhythms with either the triple dsRNA cocktail or control *gfp* dsRNA on day 0. Usually, the animals that were injected with target dsRNA showed variable behavior for 14 days after the injection (“transition” phase, day 0 - 13) before the new behavioral steady state was reached (“post” phase, day 14 – end of recording). Injection of control ds*gfp* resulted impaired or loss of circadian locomotor behavior in approximately 25% of the animals, which is comparable to previous studies (Werckenthin et al., 2020) and thus the expected loss of circadian activity based on the treatment alone.

**Figure 3:**
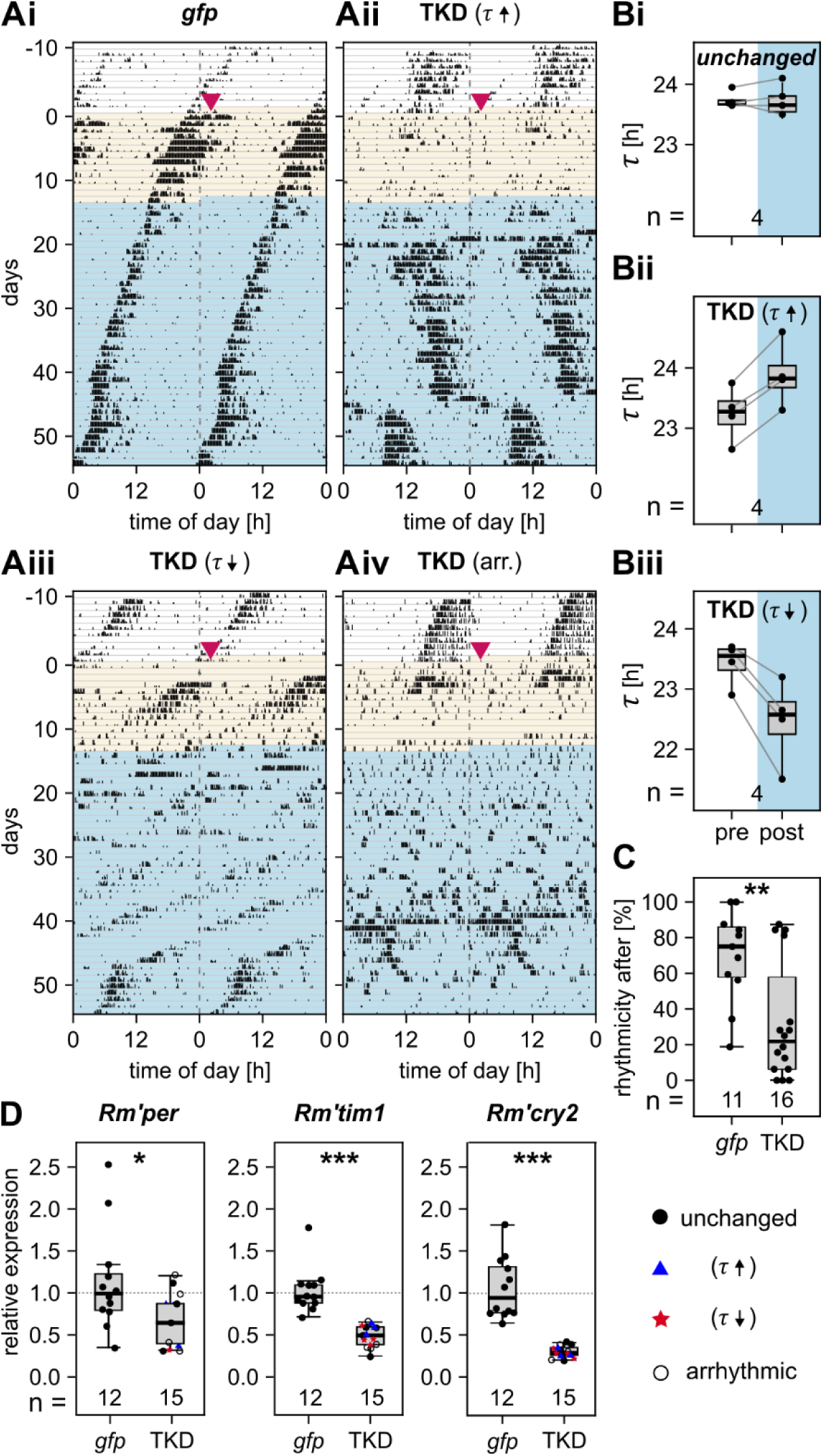
Diverse behavioral effects of RNAi triple-knockdowns (TKDs) of all negative elements of the TTFL clock in DD. **(A)** Representative double-plotted actogram of 1 control animal (*gfp*, **Ai**) and 3 example TKD animals (**Aii-iv**). **Aii** shows lengthened period (τ↑); **Aiii** shows shortened period (τ↓) and “recovery” in the last 10 days, **Aiv** shows mostly arrhythmic (arr.) behavior. Shading indicates experimental phase and is used throughout all figures: pre- (white), transition (yellow), and post-knockdown (blue); see Methods for phase definitions. Magenta arrowhead indicates day of dsRNA injection (day 0). **(B)** Illustration of the categories of period change: unchanged **(Bi)**, lengthened **(Bii)** and shortened **(Biii)**. Paired dots represent individual animals. **(C)** Rhythmicity across animals decreases in the post-phase of TKD compared to control (*gfp*) (Wilcoxon rank-sum test, W = 144, p = 0.009). **(D)** Knockdown efficiency of *Rm’per*, *Rm’tim1*, and *Rm’cry2* of TKD animals compared to ds*gfp*-injected controls, 55 days after injection at the end of the recording. Marker color and symbol indicate period change of the animal after knockdown. Statistics in Table 2.

**Table 2:** Knockdown efficiency of *Rm’per, Rm’tim1, and Rm’cry2*. Comparisons of mRNA levels of negative elements of triple-knockdown animals to dsgfp-injected controls.

| groups | test | estimate | SE | df | statistic | adj. p-value |
| --- | --- | --- | --- | --- | --- | --- |
| *Rm'per* vs. *gfp* | Welch | -0.465 | 0.193 | 15.44 | -2.408 | 0.029 * |
| *Rm'tim1* vs. *gfp* | Wilcoxon<br>rank-sum | -0.497 | N/A | N/A | 0 | <0.001 *** |
| *Rm'cry2* vs. *gfp* | Welch | -0.756 | 0.107 | 11.61 | -7.055 | <0.001 *** |

The average free-running period before dsRNA injection was τ = 23.38 ± 0.28 h for *gfp* controls (n = 11) and τ = 23.54 ± 0.37 h for TKD animals (n = 16). Control injections with *gfp* dsRNA did not significantly change the free-running period (τ_pre_ = 23.38 ± 0.28 h vs. τ_post_ = 23.50 ± 0.20 h; t(10) = 1.059, p = 0.314, n = 11). In contrast, the TKD resulted in four categories of behavior (Figure 3A, B): 1) regular, circadian activity with longer periods (Figure 3Aii, Bii; 4 of 16 animals; τ_pre_ = 23.24 ± 0.46 vs. τ_post_ = 23.90 ± 0.54 h), 2) shortened period (Figure 3Aiii, Biii; 4 of 16 animals; τ_pre_ = 23.42 ± 0.37 vs. τ_post_ = 23.3 ± 0.61 h), 3) arrhythmic behavior (Figure 3Aiv, 4 of 16 animals), and 4) unchanged (Figure 3Bi, 4 of 16; τ_pre_ = 23.75 ± 0.14 vs. τ_post_ = 23.66 ± 0.32 h). Rhythmicity analysis indicated a significant decrease in the rhythmicity percentages of the TKD animals compared to control (Wilcoxon rank-sum test gfp (n = 11) vs. TKD (n = 16), W = 144, p = 0.009, Figure 3C) but without total loss of rhythmicity based on our criteria (see Methods). In addition, knockdown efficiency for all negative elements was assessed at the end of the behavioral experiments. Independent of the behavioral category, dsRNA injection significantly decreased relative mRNA levels of each negative element in the TKDs (Figure 3D, Table 2; relative expression *Rm’per* gfp vs. TKD: 1.133±0.610 vs. 0.668± 0.308; *Rm’tim1* gfp vs. TKD: 1.027±0.271 vs. 0.487±0.129; *Rm’cry2* gfp vs. TKD: 1.054±0.366 vs. 0.298±0.068). Even with this decrease in mRNA levels, the knockdown of all known negative elements of the TTFL clockwork did not stop circadian locomotor activity to a greater extent than expected by the RNAi treatment alone.

### 6.2 Endogenous oscillations in *Rm’clk* mRNA levels

Having established that the full expression of all negative TTFL elements is not essential for maintaining circadian locomotor activity, we next investigated the role of the positive TTFL elements *Rm’clk* and *Rm’cyc* in regulating daily and circadian oscillations of behavior. Of the identified negative elements, *Rm’per*, *Rm’tim1*, and *Rm’cry2* show daily oscillations, and *Rm’tim1* and *Rm’cry2* are also expressed in a circadian manner under constant conditions (Werckenthin et al., 2012). Here, to identify daily and circadian oscillations of the positive elements *Rm’clk* and *Rm’cyc*, we used qPCR to quantify their mRNA levels in time series in LD and DD conditions, respectively, and assessed rhythmicity and cycle periods with cosinor fits.

In LD, relative expression of *Rm’cyc* significantly changed in a daily, 24 h rhythm (Figure 4Ai, Aii; Table 3), with the acrophase during the late light phase at ZT 9.73 and trough during the dark phase. Interestingly, *Rm’clk* was also periodically expressed, but in a 12 h rhythm with peaks at dusk and dawn and troughs in the middle of the day and middle of the night (Figure 4Bi, Bii; Table 3).

**Figure 4:**
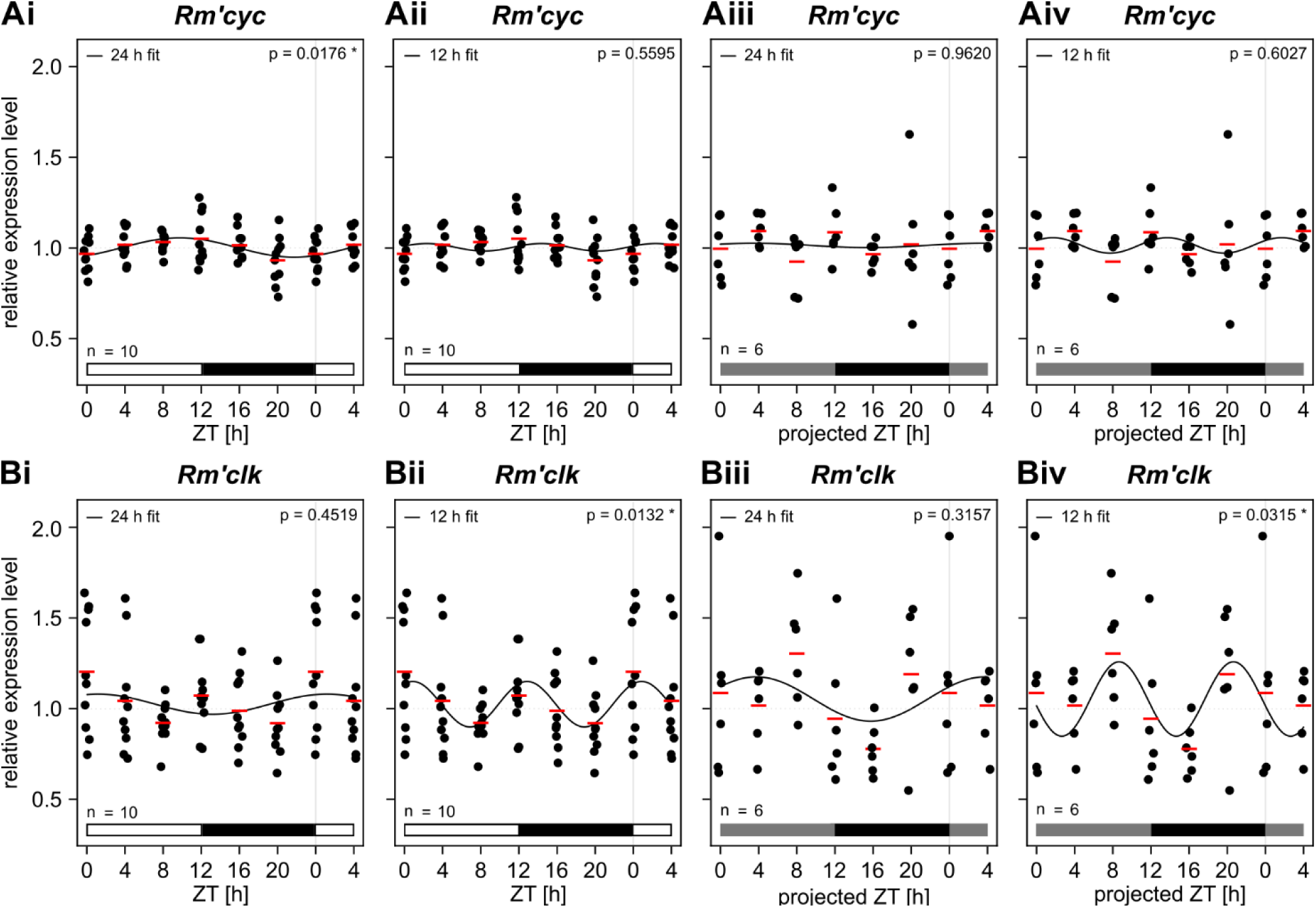
Relative mRNA expression of *Rm’cyc* and *Rm’clk* throughout the day. **(A)** Relative expression of *Rm’cyc* under 12:12 light-dark (light phase indicated by white bar, dark phase indicated by black bar) and constant darkness (DD; projected light phase indicated by grey bar) conditions. **(B)** Relative expression of *Rm’clk* under LD and DD conditions. Each panel shows: LD 24h cosinor fit (**i**), 12h cosinor fit (**ii**), DD 24h cosinor fit (**iii**), DD 12h fit (**iv**). Individual data points represent 3 pooled brains. Red horizontal bars indicate the mean. Cosinor regression parameters are summarized in Table 3. Expression values are normalized to the grand-mean of *Rm’rpl18* relative expression (2^−ΔΔCt^). Data from (projected) ZT 0 and 4 are repeated for better visualization of rhythms.

**Table 3:** Cosinor regression parameters 24h and 12h. Analysis was performed on grand-mean normalized relative expression (2−ΔΔCt, Rm’rpl18 reference) for LD (n=10 animals per ZT, 6 timepoints, N=60 total) and DD (n=6 animals per CT, 6 timepoints, N=36 total) conditions independently. N=60 (ZT), 36 (DD). Phase in ZT/CT hours from ZT0.

| target | condition | period | amplitude | acrophase | R<sup>2</sup> | p-value |
| --- | --- | --- | --- | --- | --- | --- |
| *Rm'cyc* | ZT | 24 | 0.053 | 9.73 | 0.126 | 0.018 * |
|  |  | 12 | 0.021 | 2.35 | 0.019 | 0.559 |
|  | DD | 24 | 0.012 | 3.54 | 0.002 | 0.962 |
|  |  | 12 | 0.043 | 1.7 | 0.028 | 0.603 |
| *Rm'clk* | ZT | 24 | 0.055 | 1.14 | 0.026 | 0.452 |
|  |  | 12 | 0.126 | 0.86 | 0.134 | 0.013 * |
|  | DD | 24 | 0.122 | 3.75 | 0.062 | 0.316 |
|  |  | 12 | 0.205 | 8.65 | 0.175 | 0.032 * |

In DD, circadian expression of *Rm’cyc* ceased (Figure 4Aiii, Aiv; Table 3). *Rm’clk* levels continued to oscillate with a 12 h period but acrophase shifted towards the middle of the day and middle of the night at ZT 8 and 20, respectively, compared to LD (Figure 4Biii, Biv; Table 3). Even though both are components of the circadian clock, *Rm’clk*, but not *Rm’cyc*, was endogenously oscillating.

### 6.3 Knockdown of *Rm’clk*, but not *Rm’cyc*, cancels circadian locomotor activity

After confirming endogenous oscillations of the mRNA levels of *Rm’clk*, we knocked down *Rm’cyc* and *Rm’clk* either individually or simultaneously in double knockdowns (DKD) (Figure 5), with RNAi. Similar to the TKD, we observed the locomotor activity in running wheel assays for 10 days before injection until 66 days after injection, with the same categorization as pre, transition, and post phases as described above. ds*gfp* was injected as negative control. After the injection of ds*gfp* (Figure 5Ai), the median rhythmicity across animals decreased to ∼75% (Figure 5B; Table 4) but the free-running cycle period did not change significantly (τ_pre_ = 23.38 ± 0.36 h vs. τ_post_ = 23.34 ± 0.25 h; t(42) = 0.297, p = 0.915, n = 22; Figure 5C; Table 5). In contrast, knockdown of *Rm’cyc* (Figure 5Aii) decreased the median rhythmicity across animals to ∼40% (Figure 5B; Table 4) with large interindividual variability. Where rhythmicity was maintained, the cycle period increased significantly from τ_pre_ = 23.52 ± 0.32 h to τ_post_ = 24.02 ± 0.27 h (t(26) = −4.326, p < 0.001, n = 14; Figure 5C; Table 5). However, when we knocked down *Rm’clk* (Figure 5Aiii), circadian locomotor activity was abolished in 7 of 10 animals (Figure 5B; Table 4), and even the three rhythmic animals showed only occasional bouts of circadian activity and were otherwise arrhythmic. This effect was even more pronounced in DKD animals (Figure 5Aiv), where we were unable to detect any circadian locomotor activity in any animal (Figure 5B; Table 4), demonstrating that *Rm’clk* seems to be an essential component for ongoing circadian behavior. Interestingly, both *Rm’clk* knockdowns and DKDs had increased cycle periods in the transition phase before becoming arrhythmic (n = 19; Figure 5C, D).

**Figure 5:**
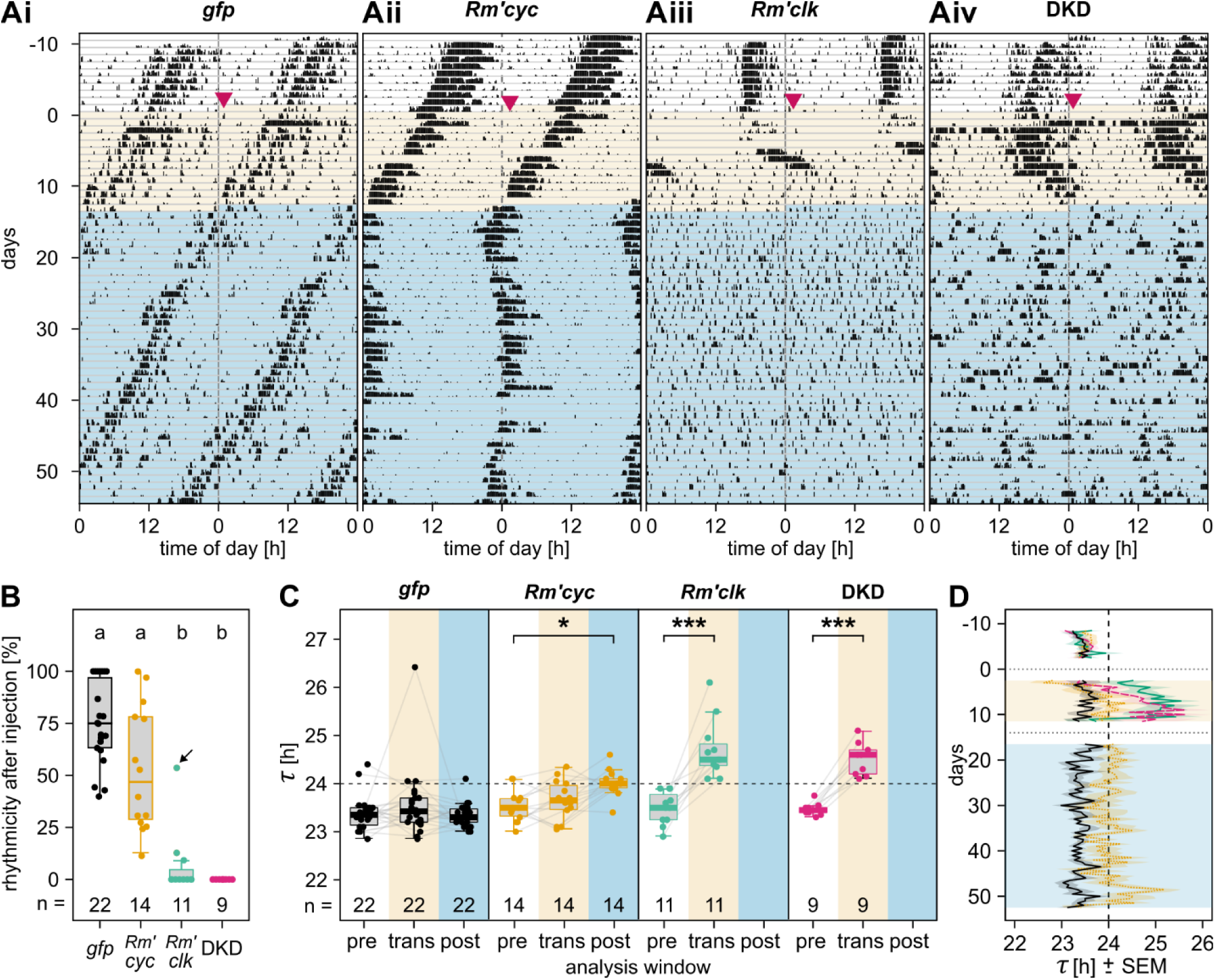
Knockdown of *Rm’clk* but not of *Rm’cyc* disrupts circadian locomotor activity. **(A)** Representative double-plotted actograms of running wheel activity recorded over 66 days for each experimental group: (**i**) *gfp* control, (**ii**) *Rm’cyc* knockdown, (**iii**) *Rm’clk* knockdown, and (**iv**) double knockdown (DKD). Magenta arrowhead indicates day of dsRNA injection (day 0). **(B)** Percentage of rhythmic windows (see Methods) for each animal (dot) during the post-injection phase per experimental group. The black arrow marks an individual that did not show clear arrhythmicity; same individual as indicated in Figure 6. Detailed statistics in Table 4. **(C)** Free-running periods (τ) estimated from sliding-window cosinor fits (see Methods) on locomotor activity recordings across the pre, trans and post phase, separately for *gfp* control, *Rm’clk*, *Rm’cyc*, and DKD animals. Data from the same animal are connected by lines. Detailed statistics in Table 5. **(D)** Trajectory of period lengths (mean ± SEM) across the duration of the recording. Horizontal lines mark phase boundaries (pre → trans → post); values are plotted at window centers, which start/stop 2.5 days after/before boundaries due to the 5-day window. Solid black: *gfp* control; dotted orange: *Rm’cyc*; solid green: *Rm’clk*; pink dashed: DKD. Window size: 5 days; step size: 0.5 days.

**Table 4:** Pairwise comparisons of rhythmicity between groups of knockdowns of positive elements of the TTFL clock. Tukey HSD post-hoc test.

| groups | estimate | SE | df | t.ratio | p-value |
| --- | --- | --- | --- | --- | --- |
| *Rm'clk* vs. DKD | 2.42 | 9.51 | 52 | 0.255 | 0.994 |
| *Rm'clk* vs. *Rm'cyc* | -50.17 | 8.88 | 52 | -5.649 | <0.001 *** |
| *Rm'clk* vs. *gfp* | -71.09 | 8.18 | 52 | -8.690 | <0.001 *** |
| DKD vs. *Rm'cyc* | -52.59 | 8.30 | 52 | -6.338 | <0.001 *** |
| DKD vs. *gfp* | -73.52 | 7.54 | 52 | -9.747 | <0.001 *** |
| *Rm'cyc* vs. *gfp* | -20.93 | 6.74 | 52 | -3.105 | 0.016 * |

**Table 5:** Pairwise comparison of period length across experimental phases. Period lengths were obtained with a sliding-window cosinor fit (see Methods). Post-phases of *Rm’clk* knockdowns and DKDs were excluded (non-significant rhythmicity). Groups with two phases (*Rm’clk*, DKD) were analyzed with paired *t*-tests. Groups with three phases (*gfp* and *Rm’cyc*) were compared with m ixed models and Tukey HSD post-hoc test.

| groups | estimate | SE | df | t.ratio | p-value |
| --- | --- | --- | --- | --- | --- |
| *gfp* (pre) vs. *gfp* (trans) | -0.191 | 0.130 | 42 | -1.468 | 0.317 |
| *gfp* (pre) vs. *gfp* (post) | 0.039 | 0.130 | 42 | 0.297 | 0.953 |
| *gfp* (trans) vs. *gfp* (post) | 0.230 | 0.130 | 42 | 1.765 | 0.194 |
| *Rm'cyc* (pre) vs. *Rm'cyc* (trans) | -0.139 | 0.114 | 26 | -1.224 | 0.450 |
| *Rm'cyc* (pre) vs. *Rm'cyc* (post) | -0.496 | 0.114 | 26 | -4.362 | <0.001 *** |
| *Rm'cyc* (trans) vs. *Rm'cyc* (post) | -0.357 | 0.114 | 26 | -3.138 | 0.011 * |
| *Rm'clk* (pre) vs. *Rm'clk* (trans) | -1.218 | 0.166 | 10 | -7.363 | < 0.001 *** |
| DKD (pre) vs. DKD (trans) | -1.044 | 0.084 | 8 | -12.506 | < 0.001 *** |

### 6.4 Decline of mRNA levels does not correlate directly with the decline of circadian locomotor activity rhythms

Since both *Rm’clk*-knockdown and DKD animals had the same qualitative behavioral phenotype, we used *clk* knockdowns to test knockdown efficiency over the course of 2 months. For this, we measured mRNA levels in whole brains at different days after the dsRNA injection (Figure 6A). Additionally, we measured the knockdown efficiency of *Rm’cyc* of the respective single-knockdown animals after 4 weeks in DD and observed as significant decreased relative mRNA levels from 1.02±0.27 in untreated animals to 0.408±0.125 (Supplementary Figure S 2).

**Figure 6:**
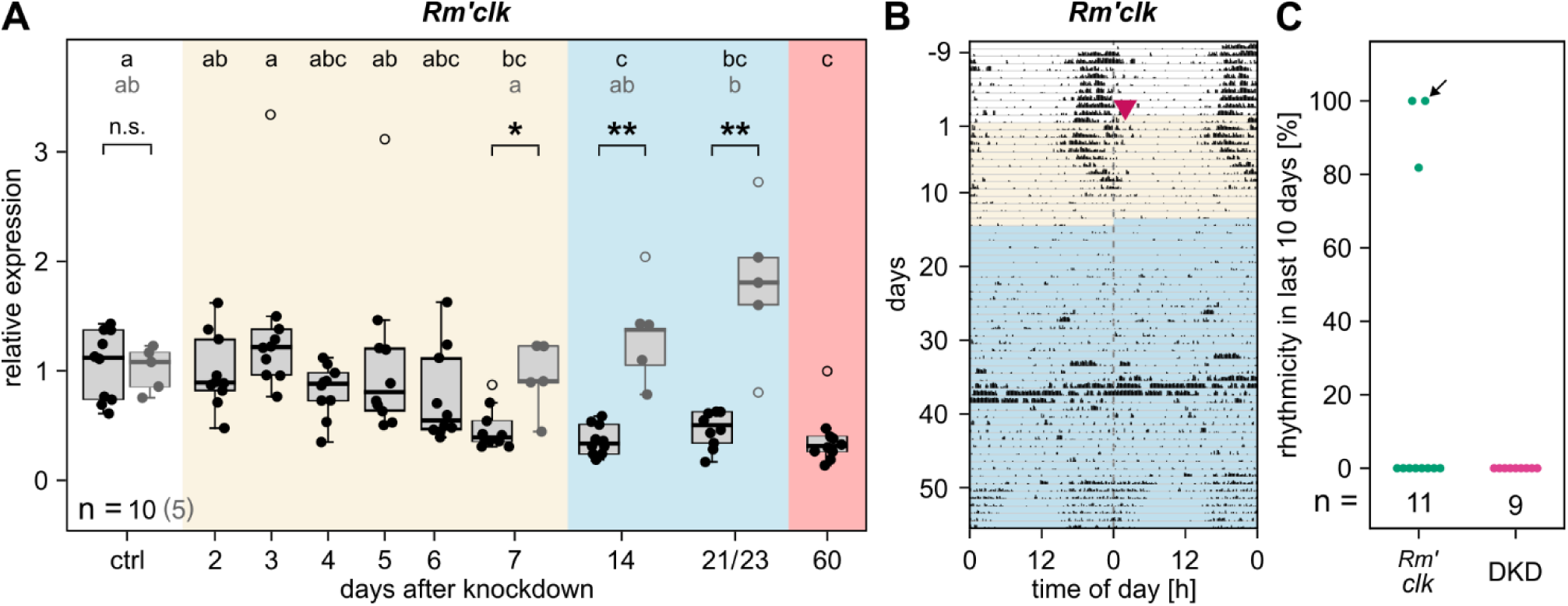
*Rm’clk* knockdown is not directly correlated to circadian locomotor rhythmicity. **(A)** Relative expression levels of *Rm’clk* mRNA reached their minimum 7 days after RNAi-dependent knockdown (black) and did not recover even after 60 days. The transition from high to low expression occurred between 4 to 7 days (Kruskal-Wallis with Dunn’s post-hoc test with Holm correction, statistics in Table 6). Relative expression levels of untreated animals (grey) did not decrease during the same time (one-way ANOVA with Tukey’s post-hoc test, Table 7). Brackets indicate statistical comparisons between matched timepoints of the knockdown and untreated animals (Wilcoxon rank-sum tests with Bonferroni correction, Table 8). **(B)** Representative double-plotted actogram of one *Rm’clk*-knockdown animal in DD that lost rhythmicity at day 11 and regained rhythmicity approximately 40 days after dsRNA injection (3 of 11 animals showed recovery). **(C)** Percentage of rhythmic windows (window-size: 5 days) over the last 10 days of *Rm’clk*-knockdown and DKD animals. Arrow indicates an *Rm’clk*-knockdown individual that did not become arrhythmic, same individual as indicated in Figure 5B.

**Table 6:** Knockdown efficiency of *Rm’clk*. Pairwise comparisons of the knockdown animals across timepoints (Kruskal-Wallis test, χ²(9) = 58.56, p < 0.001; Dunn’s post-hoc test, Holm-adjusted for 45 comparisons).

| groups | n1 | n2 | Z | p-adj |
| --- | --- | --- | --- | --- |
| ctrl - d2 | 10 | 10 | -0.293 | 1.000 |
| ctrl - d3 | 10 | 10 | 0.698 | 1.000 |
| ctrl - d4 | 10 | 10 | -1.044 | 1.000 |
| ctrl - d5 | 10 | 10 | -0.624 | 1.000 |
| ctrl - d6 | 10 | 10 | -1.734 | 1.000 |
| ctrl - d7 | 10 | 10 | -3.418 | 0.022 * |
| ctrl - d14 | 10 | 10 | -4.124 | 0.002 ** |
| ctrl - d21 | 10 | 10 | -3.237 | 0.041 * |
| ctrl - d60 | 10 | 10 | -4.143 | 0.001 ** |
| d2 - d3 | 10 | 10 | 0.990 | 1.000 |
| d2 - d4 | 10 | 10 | -0.752 | 1.000 |
| d2 - d5 | 10 | 10 | -0.331 | 1.000 |
| d2 - d6 | 10 | 10 | -1.441 | 1.000 |
| d2 - d7 | 10 | 10 | -3.125 | 0.059 |
| d2 - d14 | 10 | 10 | -3.831 | 0.005 ** |
| d2 - d21 | 10 | 10 | -2.944 | 0.097 |
| d2 - d60 | 10 | 10 | -3.850 | 0.005 ** |
| d3 - d4 | 10 | 10 | -1.742 | 1.000 |
| d3 - d5 | 10 | 10 | -1.322 | 1.000 |
| d3 - d6 | 10 | 10 | -2.432 | 0.406 |
| d3 - d7 | 10 | 10 | -4.116 | 0.002 ** |
| d3 - d14 | 10 | 10 | -4.821 | < 0.001 *** |
| d3 - d21 | 10 | 10 | -3.935 | 0.003 ** |
| d3 - d60 | 10 | 10 | -4.840 | < 0.001 *** |
| d4 - d5 | 10 | 10 | 0.420 | 1.000 |
| d4 - d6 | 10 | 10 | -0.690 | 1.000 |
| d4 - d7 | 10 | 10 | -2.374 | 0.422 |
| d4 - d14 | 10 | 10 | -3.079 | 0.064 |
| d4 - d21 | 10 | 10 | -2.193 | 0.651 |
| d4 - d60 | 10 | 10 | -3.098 | 0.062 |
| d5 - d6 | 10 | 10 | -1.110 | 1.000 |
| d5 - d7 | 10 | 10 | -2.794 | 0.151 |
| d5 - d14 | 10 | 10 | -3.499 | 0.017 * |
| d5 - d21 | 10 | 10 | -2.613 | 0.251 |
| d5 - d60 | 10 | 10 | -3.519 | 0.016 * |
| d6 - d7 | 10 | 10 | -1.684 | 1.000 |
| d6 - d14 | 10 | 10 | -2.389 | 0.422 |
| d6 - d21 | 10 | 10 | -1.503 | 1.000 |
| d6 - d60 | 10 | 10 | -2.409 | 0.416 |
| d7 - d14 | 10 | 10 | -0.705 | 1.000 |
| d7 - d21 | 10 | 10 | 0.181 | 1.000 |
| d7 - d60 | 10 | 10 | -0.725 | 1.000 |
| d14 - d21 | 10 | 10 | 0.886 | 1.000 |
| d14 - d60 | 10 | 10 | -0.019 | 1.000 |
| d21 - d60 | 10 | 10 | -0.906 | 1.000 |

**Table 7:** Expression of *Rm’clk* in untreated animals. Pairwise comparisons of the untreated animals across timepoints. One-way ANOVA, F(3,16) = 3.57, p = 0.038; estimated marginal means with Tukey-adjusted p-values.

| groups | estimate | SE | df | t.ratio | p-adj |
| --- | --- | --- | --- | --- | --- |
| ctrl - d7 | 0.078 | 0.291 | 16 | 0.267 | 0.993 |
| ctrl - d14 | -0.338 | 0.291 | 16 | -1.159 | 0.660 |
| ctrl - d23 | -0.777 | 0.291 | 16 | -2.667 | 0.072 |
| d7 - d14 | -0.415 | 0.291 | 16 | -1.426 | 0.502 |
| d7 - d23 | -0.855 | 0.291 | 16 | -2.934 | 0.043 * |
| d14 - d23 | -0.439 | 0.291 | 16 | -1.508 | 0.455 |

**Table 8:** Comparison of *Rm’clk* expression between knockdown and untreated animals. Wilcoxon rank-sum test with Bonferroni adjustment for four comparisons.

| groups<br>(knockdown vs.<br>untreated) | n1 | n2 | U | p-adj |
| --- | --- | --- | --- | --- |
| ctrl - ctrl | 10 | 5 | 27.00 | 1.000 |
| d7 – d7 | 10 | 5 | 3.00 | 0.034 * |
| d14 – d14 | 10 | 5 | 0.00 | 0.011 * |
| d21 – d23 | 10 | 5 | 0.00 | 0.011 * |

The dsRNA injection did not immediately reduce *Rm’clk* mRNA levels. Only on day 7 after injection, relative expression decreased significantly from 1.05 ± 0.32 to 0.47 ± 0.19, remaining at minimum for at least 60 days after injection (Figure 6A black data points, Table 6). This was not an aging effect in the long-lived (∼2 years) cockroaches since the respective mRNA levels did not decline in untreated animals in the same time window (Figure 6A grey data points, Table 7, Table 8). This time course of mRNA decline did not correlate directly with the time course of decline in circadian locomotor behavior. Period lengthening started in the transition phase on day 3 after injection, when mRNA levels were still at control values (cf., Figure 5D, Figure 6A). Furthermore, the new behavioral steady state was only reached 14 days after injection, while mRNA levels already reached effective knockdown after 7 days. In addition, a few (2 of 11) animals recovered circadian locomotor activity approximately 50 days after injection (τ = 22.85 and 23.6; Figure 6B). However, *Rm’clk* mRNA levels were still minimal after 60 days, indicating recovery mechanisms different form upregulation of the expression of this gene. This demonstrates that changes in circadian locomotor activity did not correlate directly with changes in *Rm’clk* mRNA levels.

### 6.5 Heterarchical model architecture adequately recapitulates the biological findings

So far, our results indicate that the circadian locomotor activity is partially independent of a functional TTFL clockwork. To explore the possibility and extent to which a heterarchical clock network, that consists of independent but interacting TTFL and PTFL clocks, could account for our experimental results, we constructed a computational model of the circadian pacemaker network of the cockroach in constant conditions. TTFL and PTFL oscillators were modeled by individual PSAS. One such TTFL and PTFL oscillator were mutually coupled to construct a morning oscillator (MO) cell with a cycle period <24 h; another TTFL and PTFL oscillator were mutually coupled to construct an evening oscillator (EO) cell with a cycle period >24 h, reminiscent of the model of morning and evening oscillators as demonstrated in, e.g., *Drosophila* (Yoshii et al., 2012). Based on previous knockdown experiments in the Madeira cockroach (Werckenthin et al., 2020), the putative MO and EO neurons comprise different negative elements of the TTFL clock, respectively. Therefore, our model MO-TTFL contains interacting *clk* and *per,* and the EO-TTFL interacting *clk* and *tim1*. In addition to the TTFL clock, each modeled MO and EO contained a PTFL clock. The PTFL clocks control circadian release of neuropeptides as coupling factors between the MO and EO cells. Therefore, the PTFL clocks are tightly linked to circadian oscillations in membrane potential, intracellular Ca^2+^ concentration and cAMP levels (Rojas et al., 2019; Stengl et al., 2015; Stengl and Arendt, 2016; Stengl and Schneider, 2024; Wei et al., 2014; Wei and Stengl, 2012).

The oscillations of the obtained PSASs accurately approximated the periods of the TTFLs and PTFLs, corresponding to the behavioral periods found in the biological experiments (c.f. Figure 2D, Figure 5C). In the uncoupled case, the PSAS oscillator modeling the EO-TTFL exhibited a period of 1.0147 days (24.35 h), the PSAS corresponding to the MO-TTFL had a period of 1.0032 days (24.08, the EO-PTFL had a period of 1.0514 days (25.23 h), and the MO-PTFL one of 0.9325 days (22.38 h). Prior to knockdown, the connections between TTFL and PTFL oscillators, as well as those among PTFL oscillators, were governed by constant coupling gains *k̅_ij_*, consistent with established studies (Rosenblum et al., 2001; Schmal et al., 2018; Wickramasinghe and Kiss, 2013) which investigate both intercellular and intracellular coupling using fixed coupling strengths.

For the heterarchical network, the coupling gains were selected to:

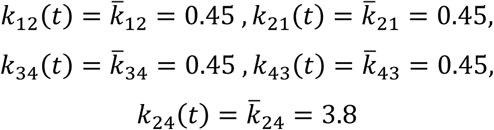

for *t* ∈ [*t*_0_, *t_k_*]. Note that that *k*_12_(*t*) = *k̅*_12_ and *k*_34_(*t*) = *k̅*_34_ for all *t* ∈ [*t*_0_, ∞[. The gains were set to the same value, except for *k̅*_24_, to reflect the mutually interacting nature of the heterarchical network. This configuration ensured that both TTFL and PTFL oscillators contributed equally to the “behavior” of the “cockroach”. Before the knockdown, the gains *k̅*_21_ and *k̅*_43_ cannot be chosen larger than above to avoid periods larger than 1 day, which would contradict our experimental findings. The gains *k*_12_(*t*) = *k̅*_12_ and *k*_34_(*t*) = *k̅*_34_ were also chosen with the same magnitude, otherwise oscillations with larger amplitudes would be induced for the TTFLs after the knockdown due to the input from the oscillating PTFLs. However, *k̅*_12_ and *k̅*_34_ cannot be chosen arbitrarily small since this would incur oscillator behavior which does not comply with the biological measurements.

By introducing the additional gains 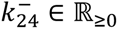 and 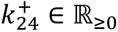, the time-varying coupling gain between the two PTFLs is defined to:

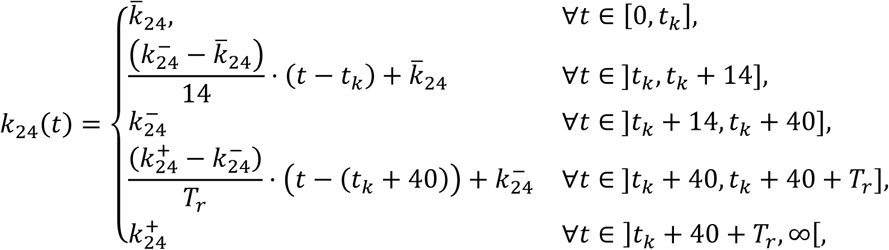

where *T_r_* ∈ ℝ_>0_ represents the duration of the recovery phase in which the behavioral rhythm returned in the biological experiments. Assuming that the link between the PTFL oscillators diminishes following knockdown, the weight of this interaction (represented by *k*_24_(*t*)) is gradually reduced after *t_k_* and remains near zero until *t* = *t_k_* + 40. During recovery, the weight increases linearly, rising to a final value of 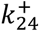. The link from EO-PTFL to MO-PTFL is excluded, thus *k*_42_(*t*) = 0, since we hypothesized that the MO cell dominates the behavior in DD. To capture the increasing period for *t* ∈]*t_k_*, *t_k_* + 7], the coupling gain *k*_21_(*t*) is temporarily increased to 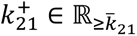 over this interval as follows:

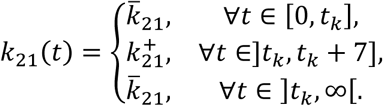

Similarly, *k*_43_(*t*) is defined using 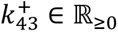, following the same structure as for the previous coupling gain. In addition, the parameters 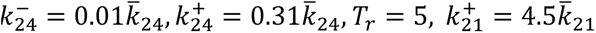, 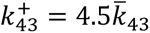 and the knockdown effect *u_i_*(*t*) were chosen to reproduce the experimentally observed behavior in the networked system (Eq. (1); Figure 7).

**Figure 7:**
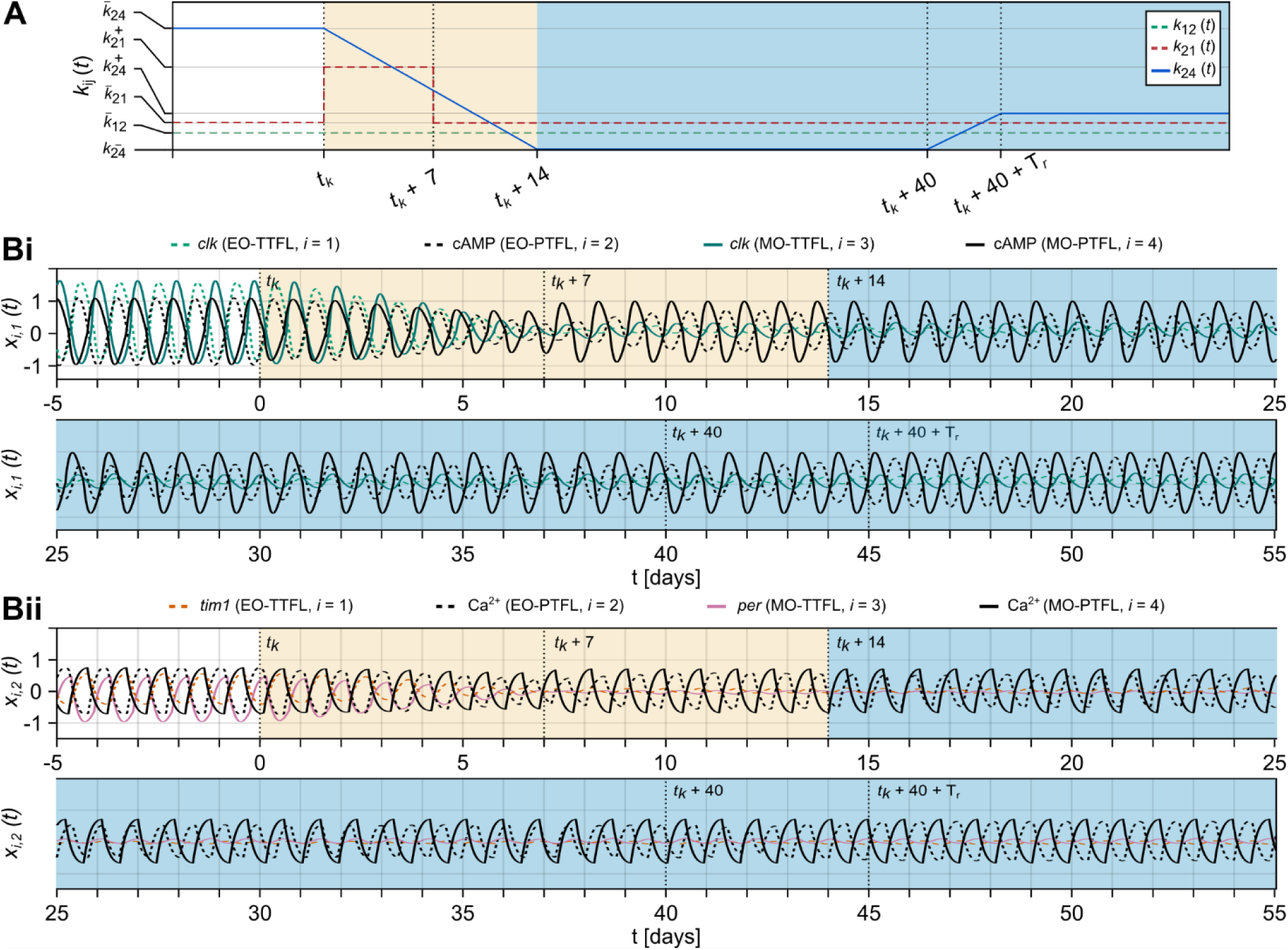
Simulation of the heterarchical model of coupled TTFL and PTFL PSAS oscillators in DD. **(A)** Change in coupling gains over simulation time. Colors are the same as in Figure 2A. **(B)** Simulations of Eq. (1) for *x_i_*_,1_(*t*) (**i**) and *x_i_*_,2_(*t*) (**ii**) for *i* ∈ {1,2,3,4}. Bottom plots are continuations of the top plots. The coupling gains are chosen such that the network recapitulates the behavior observed in the biological experiments. The vertical lines mark the separation into the five intervals referred to in the text where the coupling gains changed.

We used the model to simulate the knockdown of *clk*, which resulted in arrhythmic behavior in most cockroaches and subsequent recovery in a subset, both of which did not strictly correlate with *clk* mRNA levels (Figure 7). The model is assumed to represent rhythmic behavior of the insect if the MO- and EO-PTFLs oscillate in antiphase. The modeling results can be broken down into five time intervals: 1) *t*_0_ to *t_k_*, which represents the “wild-type” steady state until the initiation of *clk* knockdown at *t_k_* (pre-phase of the biological experiments); 2) *t_k_* to *t_k_* + 7, which represents the first 7 days after knockdown initiation in which *clk* mRNA levels steadily decrease to their minimum (first half of trans-phase in the biological experiments); 3) *t_k_* + 7 to *t_k_* + 14, in which the behavior reaches a new steady state (second half of the trans-phase in the biological experiments); 4) *t_k_* + 14 to *t_k_* + 40, which represents the steady state of arrhythmic activity (post-phase in the biological experiments); 5) *t_k_* + 40 to the end of the simulation, in which the rhythmic behavior recovers.

In the first interval, oscillations corresponded to rhythmic behavior since both PTFLs were in anti-phase with a period of approximately 0.95 days (c.f., Figure 5C). Further simulations show that the period keeps increasing to values larger than 1 day for increasing coupling gains *k̅*_21_ and *k̅*_43_. At the beginning of the second interval, the knockdown is initiated. Subsequently, the amplitude of the TTFLs gradually decreases (c.f., Figure 6A). The period increased up to values of about 1.05 days (c.f., Figure 5C, D) and the phase lags between the oscillators started to shift. During the third interval, the coupling gain *k*_24_(*t*) was gradually reduced to 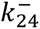. A smaller value here would make the behavior already arrhythmic, i.e., the PTFLs would lose their antiphase relationship. In the fourth interval, where the coupling gain *k*_24_(*t*) was reduced to 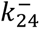, the PTFLs did not remain in anti-phase or expressed any constant phase lags but rather oscillated at individual periods, thus representing arrhythmic behavior (c.f., Figure 5). At the beginning of the fifth interval, the coupling gain *k*_24_(*t*) was gradually increased to 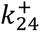, leading to emerging rhythmic behavior, i.e., the PTFLs regained their antiphase oscillations, however, the TTFLs were still affected by the knockdown and did not recover to their initial state (c.f., Figure 6). Overall, the heterarchical network describes all biological findings apart from the lengthened period after the knockdown for *t* ∈]*t_k_*, *t_k_* + 14] (only occurring up to approximately *t_k_* + 4.5).

### 6.6 Hierarchical model architecture fails to recapitulate behavioral observations

In contrast to the heterarchical network, the PTFLs of the hierarchical network do not constitute oscillating systems but are modeled by transient systems. Here, one can distinguish two different cases: The Hurwitz Matrix *A* has 1) complex conjugate eigenvalues and the uncoupled PTFLs exhibit a damped oscillations towards the steady state 0_2_; 2) real eigenvalues and the states of the uncoupled PTFLs asymptotically converge to the steady state 0_2_.

Considering the first case, a matrix *A* with complex eigenvalues *λ*_1,2_ = −3 ± 3*j* was chosen. While the time evolution of the coupling gains (Figure 7A) was chosen to be the same as for the heterarchical network, the constant coupling gains were chosen as

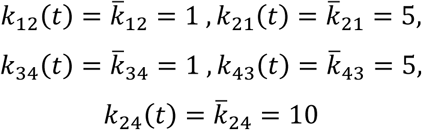

for *t* ∈ [*t*_0_, *t_k_*]. In the simulation of this hierarchical network (Figure 8), the PTFLs synchronized before knockdown in antiphase with period *T* ≈ 0.96 days, which corresponds matches the output of the heterarchical network before knockdown.

**Figure 8:**
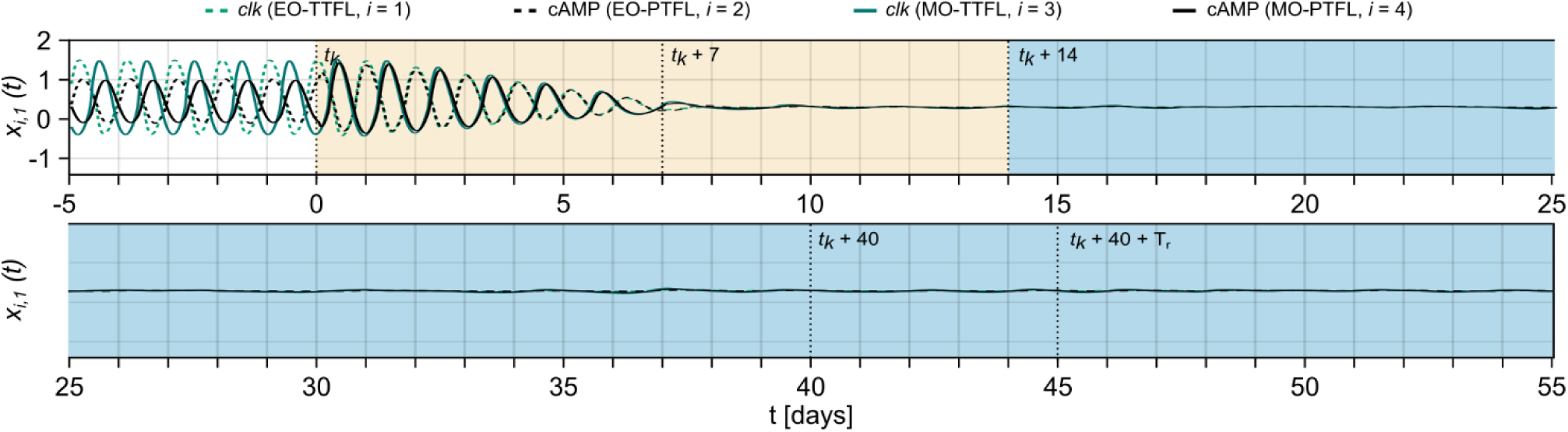
Simulation of the hierarchical network. Simulation of the hierarchical network showing only *x_i_*_,1_(*t*) for *i* ∈ {1,2,3,4} for simplification. The *A* matrix of the dynamics of the transient PTFL has complex eigenvalues and the coupling gains are adapted to the hierarchical network structure. Relative changes in coupling gains are the same as in Figure 7.

For the second case, a matrix *A* with real eigenvalues *λ*_1_ = −2 and *λ*_2_ = −1 was chosen, together with constant coupling gains

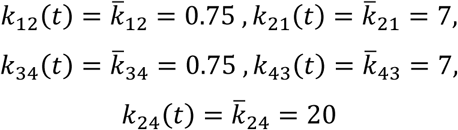

for *t* ∈ [*t*_0_, *t_k_*]. The corresponding simulation (Supplementary Figure S 3) shows that the PTFLs also synchronize with a constant phase relation and period *T* ≈ 0.95 before knockdown. At *t* = *t_k_* + 7 the knockdown deleted periodic oscillations of the TTFLs in both cases which ceased rhythmic output of the PTFLs at that time, which would correspond to loss locomotor activity in the cockroach. Even after *t* = *t_k_* + 40 the PTFLs do not regain their rhythmic behavior as it was the case for the heterarchical network. Thus, the hierarchical model fails to fully describe cockroach behavior after knockdown.

### 6.7 Model prediction of cockroach behavior in constant light conditions after knockdown

Deriving a computational model also allows us to make predictions of what may happen under different circumstances. As this model was designed based on the circadian clock in DD, simulating the case of constant light (LL) is an obvious choice. In LL, the free-running period of locomotor activity in cockroaches is longer than in DD, presumably because the dawn-locked, slower MO is now the dominant oscillator compared to DD conditions (Barrett and Page, 1989; Pittendrigh and Daan, 1976; Werckenthin et al., 2020). We had to minimally adjust the model to recapitulate the reported increase in the free-running period of locomotor activity (Barrett and Page, 1989): 1) the coupling from MO-PTFL to EO-PTFL with coupling gain *k*_24_(*t*) is reversed to a coupling from EO-PTFL to MO-PTFL with coupling gain *k*_42_(*t*), taking the same values as the previous *k*_24_(*t*), while the rest of the coupling gains *k_ij_*(*t*) stay unchanged; 2) the delay now takes the value of half the period of the dominating EO-PTFL, i.e, *t_D_*_,42_ = 1.0514/2. In the LL simulation (Figure 9A), the PTFLs are synchronized in antiphase with period *T* ≈ 1.04 days before the knockdown. After knockdown initiation, around *t* = *t_k_* + 7 the antiphase relationships between MO and EO cells were lost, but the MO and EO cells briefly regained a constant phase relationship before *t* ≈ *t_k_* + 14, which was lost shortly afterwards. Similar to DD, the PTFLs synchronized again with a constant but not entirely anti-phase phase shift some time after *t* = *t_k_* + 40 with a shortened period of *T* ≈ 0.79 days, indicating the possibility of recovery. In contrast we observed no increase in period during the transition phase as we did for the DD model and in the experimental data.

**Figure 9:**
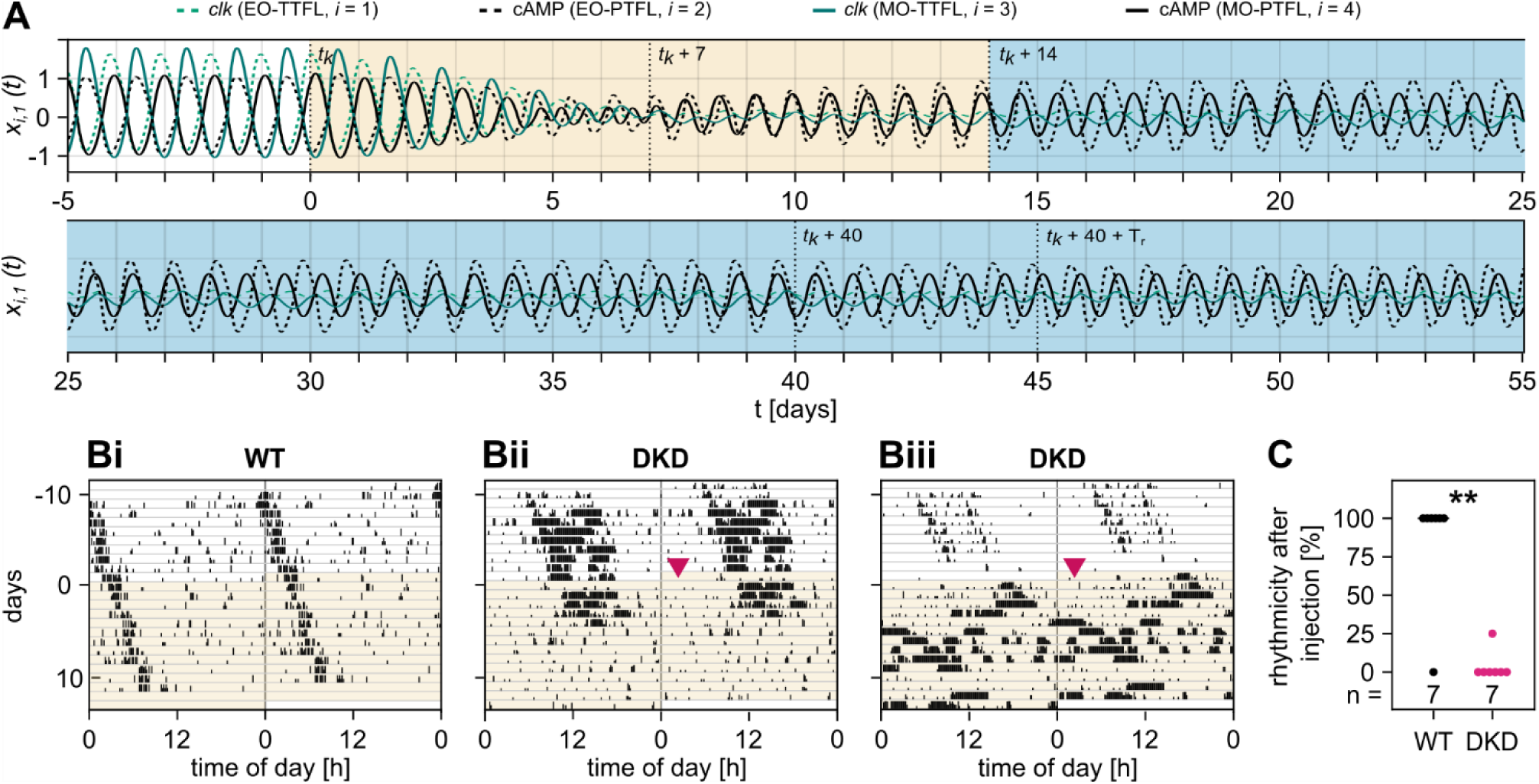
Model evaluation in constant light (LL) conditions. **(A)** Simulation of the heterarchical network after adjustment to represent constant light conditions. For simplification, only *x_i_*_,1_(*t*) for *i* ∈ {1,2,3,4} are shown. TTFL disruption including coupling gain changes are the same as in Figure 7. **(B)** Three representative double-plotted actograms of running wheel activity recorded over 24 days in LL: (**Bi**) untreated wild-type (WT) control, (**Bii -iii**) double knockdown (DKD) of *Rm’cyc* and *Rm’clk*. Magenta arrowhead indicates day of dsRNA injection (day 0). The animal in Bii retained rhythmicity in 25% of analyzed windows. **(C)** Percentage of rhythmic windows (see Methods) for each animal (dot) during the trans-phase per experimental group. Rhythmicity was significantly lower in DKD than in WT control animals (Wilcoxon rank-sum test, W = 45, p = 0.005). 6 of 7 WT controls stayed rhythmic while 6 of 7 DKD animals became fully arrhythmic and one retaining rhythmicity at 25%.

To evaluate the LL simulation predictions in biological experiments, we performed running wheel assays in LL with DKD after 10 days. Qualitatively, the biological experiments matched the model prediction (Figure 9B, C). Before knockdown, the free-running period of DKD animals in constant light was significantly longer than that of their DD counterparts (LL: τ_pre_ = 24.09 ± 0.25 h, n = 7; DD: τ_pre_ = 23.46 ± 0.17 h, n = 9; Welch’s t-test, t(10.0) = 5.79, p < 0.001). The free-running period of untreated wild-type (WT) controls (τ_pre_ = 24.34 ± 0.24 h, n = 7) and DKD animals did not differ in the pre-phase (Welch’s t-test, t(11.9) = 1.96, p = 0.074). As predicted by the heterarchical LL model, and in contrast to DD, DKD animals already became arrhythmic (6 of 7 animals) early in the trans-phase, indicating that the model captured key aspects of the interconnected TTFL and PTFL clocks. WT controls continued to express rhythmic circadian locomotor behavior (6 of 7 animals) during the same time window.

## 7 Discussion

Previous attempts at disrupting the circadian locomotor behavior of the Madeira cockroach by individually targeting the negative elements of the TTFL clock with RNAi resulted in a decrease of the free-running circadian periods of locomotor activity (Werckenthin et al., 2020). Here, we demonstrated for the first time that, of all the known positive and negative elements of the cockroach TTFL clockwork, solely the knockdown of the positive element *Rm’clk* alone was sufficient to disrupt circadian locomotor activity in DD for several weeks. Even triple knockdowns of all negative elements did not cancel circadian behavior, hinting at an additional, still unknown negative element or negative feedback mechanism that stabilizes the cockroach TTFL clock, such as possibly *clockwork orange* (Matsumoto et al., 2007).

Unexpectedly, however, we did not find a direct correlation between the decline of *Rm’clk* mRNA levels and the decline of circadian locomotor activity rhythms. In contrast to previous interpretations in different species (e.g., *tim*, *clk*, or *vri* knockdown in *Drosophila*: (Matsumoto et al., 2007)), where RNAi-dependent knockdowns did not result in complete and correlated arrhythmicity, here, we suggest a different hypothesis than mosaic or variable knockdown of mRNA levels. Instead, we hypothesize that compensatory PTFL clocks are responsible for remaining or regained circadian rhythms in behavior. Accordingly, our experimental findings could be adequately simulated with a computational model with heterarchical network architecture (systemic model), in which oscillations and coupling of PTFLs alone, without TTFL contribution, could produce an output that corresponds to circadian behavioral activity. This indicates that PTFL clocks could be a main element of a compensatory, stabilizing mechanism in circadian timing in the Madeira cockroach.

### 7.1 In different species, either *clk* or *cyc* express endogenous circadian oscillations, possibly facilitating differential phase-control of MO and EO clocks

In contrast to daily LD cycles, where both transcription factor genes *Rm’clk* and *Rm’cyc* were rhythmically expressed, only the expression rhythm of *Rm’clk* persisted in DD, with similarly low cycle amplitudes as reported previously for the cycling of *Rm’per*, *Rm’tim1*, and *Rm’cry2* (Werckenthin et al., 2012). Thus, in the Madeira cockroach, as in *Drosophila*, only *clk* but not *cyc* is rhythmically expressed. In *Drosophila*, CYC is subjected to 24 h cycles through stabilization by cycling CLK levels (Bae et al., 2000; Liu et al., 2017; Rutila et al., 1998). The cycling of *clk* but not *cyc*, even in LD cycles, is also common in many other insect species, e.g., cricket (Shao et al., 2008), sawfly (Bembenek et al., 2007), or silkworm (Sehadová et al., 2004). While homozygous *Drosophila cyc* mutants show arrhythmic behavior, the period of circadian locomotor activity is increased in heterozygous mutants as compared to wild-type flies, as expected for lower expression levels of *cyc* (Rutila et al., 1998). In contrast, *cyc* is periodically expressed in brains of honeybees and mosquitos in both LD and DD, with transcript abundance peaking at dawn and the trough occurring at dusk (Gentile et al., 2009; Rubin et al., 2006). In sandflies, expression is slightly shifted so that *cyc* peaks during the day (Meireles-Filho et al., 2006). In rat and mouse, expression of *cyc* homologue *Bmal1* oscillates with a 24 h period in the suprachiasmatic nucleus in both LD and DD with peak mRNA levels during the night, while its dimerization partner *clk* is constitutively expressed (Honma et al., 1998; Maywood et al., 2003; Taguchi et al., 2001). Apparently, the respective cycling transcription factor keeps a functional nuclear localization sequence (NLS) while the other just needs to keep protein-protein binding domains to be able to form heterodimers. In *Drosophila*, CLK contains one functional NLS, whereas CYC is lacking an NLS but nevertheless promotes the nuclear localization of CLK via heteromerization with CLK (Hung et al., 2009). In mammals, the respective roles are reversed: Although mammalian CLK does not have a functional NLS, it appears to depend on heteromerization with BMAL1, which contains one functional NLS, for nuclear translocation (Kwon et al., 2006). Thus, based on our experiments, we hypothesize that Rm’CLK but not Rm’CYC contains an NLS.

Suppression of *Rm’cyc* led to an increase in the free-running cycle period of locomotor behavior but did not result in arrhythmic animals, as is the case in *Drosophila* (Rutila et al., 1998). It is possible that the residual CYC protein levels were still sufficient to form functional CLK-CYC heterodimers to keep the TTFL running, but with a longer cycle period due to decreased abundance of heterodimers (Leloup and Goldbeter, 2003). Alternatively, we cannot exclude redundancy of heteromerization partners for Rm’CLK in the cockroach, although it has not been reported in other species. Knockdown of *Rm’clk* had a more striking effect on behavioral rhythmicity, effectively disabling the TTFL in most cockroaches. This effect was enhanced in the DKDs that lacked both positive elements, corroborating the synergistic effects of the two proteins.

While the expression patterns of *cyc/Bmal1* and *clk* differ between species, the negative elements *per*, *cry2*, and *tim* oscillate at the same phase in cockroach, honeybee, and mosquito (Gentile et al., 2009; Rubin et al., 2006; Werckenthin et al., 2012). Therefore, phases and robustness of the expression rhythms of the negative elements of the TTFL clock appear to be more conserved during evolution with more robust phase-synchronization to dusk and dawn of the LD cycle, than the expression of the positive elements with their spread of phase-relationships. Apparently, both positive elements cycled earlier in evolution and, possibly, it became advantageous that one of them lost endogenous LD-entrained cycling. This may have facilitated for different circadian clock neuron populations to cycle at distinctly different phases such as the antiphasic MO and EO clocks that control the biphasic activity pattern in *Drosophila* (Grima et al., 2004; Helfrich-Förster, 2001; Pittendrigh and Daan, 1976; Stoleru et al., 2004). Accordingly, in whole-head extracts of *Drosophila*, *clk* cycles with bimodal peaks at ZT 5 and ZT 23, reminiscent of antiphasic MO and EO clocks (Darlington et al., 1998). Also, in our study, we used whole brain extracts for qPCR, which would contain both the MO and EO clock circuits per AME. If both clock circuits express *Rm’clk* at similar levels, their antiphase oscillations could result in two expression peaks, explaining the 12 h cycles of *Rm’clk* found here.

### 7.2 No temporal correlation of mRNA levels of the TTFL clockwork and circadian locomotor activity rhythms

Unexpectedly, and in contrast to the hierarchical TTFL hypothesis, we did not observe a direct temporal correlation between RNAi-dependently declining mRNA levels and a simultaneous decline in behavioral rhythmicity. This could be due to variable mRNA depletion in distinct clock cell populations, while we could exclude age-dependent effects of mRNA decline in age-matched experimental animals. While cockroach species are very susceptible to RNAi, even much more than *Drosophila* (Bellés, 2010; Garbutt et al., 2013), we do not have the tools available to examine whether RNAi knockdown is occurring simultaneously, equally, and completely in all clock neurons. The RNAi knockdown of specific targets reduces the respective mRNA levels to a certain extent but does not necessarily completely deplete it. Only partial loss of respective mRNA levels in mosaic expression was hypothesized to be responsible for a fraction of treated animals in different species that retained daily/circadian rhythms in behavior (e.g., *tim*, *clk*, or *vri* knockdown in *Drosophila*: (Matsumoto et al., 2007), *tim* knockdown in *L. striatellus:* (Jiang et al., 2018); *cry2* knockdown in *G. bimaculatus:* (Tokuoka et al., 2017)). Without available tools for *Rhyparobia*, such as TTFL clock protein specific antisera, we could not exclude that our RNAi knockdowns resulted in mosaic expression of clock genes across tissues, or across cells within a tissue, and, thus, had to rely on circadian locomotor behavior as output of the circadian clock. Nevertheless, RNAi reduced the mRNA levels of all clock genes significantly in whole brain extracts. While the injection of any dsRNA resulted in arrhythmic locomotor behavior in approximately 20-30 % of the treated animals (this study and e.g., (Werckenthin et al., 2020)), injection of *Rm’clk* and/or *Rm’cyc* canceled circadian rhythmicity in significantly more animals than would have been expected from the effect of the injection itself.

Furthermore, as expected for a decrease in the activation of transcription of the negative elements in clock neurons controlling behavioral rhythms, knockdown of either only the single positive element *Rm’clk* or of *Rm’clk* and *Rm’cyc* together (DKD), both increased the period of circadian locomotor activity in the transition-phase. Accordingly, knockdown of the negative elements, resulting in faster or stronger activation of transcription, shortened the cycle period (Werckenthin et al., 2020). Thus, our RNAi-dependent knockdown of TTFL core genes targeted the neuronal clock neuron populations that control behavioral rhythms already in the transition phase, making mosaic-effects unlikely.

Generally, RNAi in insects results in a slow (1-2 weeks to maximum knockdown effect) but long-lasting (> months-long) reduction of target mRNA levels (Uryu et al., 2013). Even though posttranscriptional buffering of mRNA in general can last for hours to days in some insects (Becker et al., 2018; Nirmala et al., 2005), dsRNA remains for at least 24 h in the cockroach hemolymph (Garbutt et al., 2013), indicating that RNAi outlasts mRNA buffering. In *Drosophila*, peak and trough protein levels of Per and Tim lag approximately 6 hours behind peak transcription and peak mRNA levels (So and Rosbash, 1997). The *per* mRNA levels are under posttranscriptional control and thereby lag during the rising phase of transcription, indicating a more stable half-life. During the decreasing phase they match transcription rate, which indicates a less stable half-life for the mRNA during decline. Computational modeling has shown that long half-lives (> 10 h) lead to a loss of detectable circadian oscillations (Lück et al., 2014). Since we observed recovery of rhythmic locomotor activity in some *Rm’clk* knockdown animals after several weeks when mRNA levels were still at their RNAi-dependent minimum, it is unlikely that this effect was due to TTFL recovery, even when considering mRNA buffering.

The oscillations of *Rm’clk* mRNA levels in LD and DD conditions suggest that this gene plays critical role in regulating circadian locomotor activity in the Madeira cockroach. Accordingly, knocking down *Rm’clk*, but not *Rm’cyc*, resulted in the loss of rhythmic locomotor behavior in the majority of the observed cockroaches. This effect was enhanced in DKDs that lacked both respective proteins, consistent with a presumed heteromerization of both transcription factors.

It is established for *Drosophila* that the transcription factors CLK and CYC form heterodimers that translocate to the nucleus and activate transcription of the negative elements (Darlington et al., 1998; Huang et al., 2012; Liu et al., 2017; Rutila et al., 1998). While their turnover depends, among others, on phosphorylation states, partner binding and localization, degradation is on the order of hours to complete and maintain the circadian cycle (Hung et al., 2009; Kwon et al., 2006; Liu et al., 2017). Because no specific antisera against cockroach CLK are available, and antisera from other species were not successful in Western blots, we do not know yet how fast CLK protein levels decline in the cockroach. However, given the evolutionary conservation of clock systems, we assume clock proteins are degraded on a similar timescale in the cockroach as in *Drosophila*. Therefore, once our RNAi effectively reduced mRNA levels, protein levels should correspondingly decline within a day. Effective nuclear localization requires the dimerization of CLK and BMAL1/CYC (Hung et al., 2009; Kwon et al., 2006). Thus, the increase in the free-running circadian period that we observed in the transition phase of both single knockdowns might reflect this impairment in nuclear translocation. This aligns with general oscillator theory, e.g., the Goodwin model extended by Leloup and Goldbeter (2003), which predicts that a lower rate of *Bmal1* transcription leads to a more delayed inhibition of transcription, resulting in an increase in period.

Our results are not consistent with the hierarchical TTFL clock hypothesis, since knockdown manipulations did affect overall circadian locomotor behavior, while the mRNA levels of the TTFL clock did not strictly correlate with the changes in behavioral output in DD. The disruption of circadian locomotor rhythms immediately after the dsRNA injection was most likely a stress reaction due to the injection procedure and animal handling. The mRNA levels reached their minimum on day 7 after dsRNA injection, and therefore, the respective protein levels were presumably minimized shortly after that. However, the increase in cycle period of the locomotor activity rhythms continued in the following days, up to reaching arrhythmicity depending on the type of knockdown. Most inconsistent with the TTFL hypothesis of circadian timing in the Madeira cockroach is that some cockroaches regained behavioral rhythmicity in DD approximately 50 days after injection while mRNA levels were still at their minimum.

### 7.3 Our novel hypothesis states that PTFL mechanisms explain timing in the Madeira cockroach

Taken together, our results provide evidence for the existence of additional, compensatory posttranslational mechanisms with feedback loop control besides the canonical circadian TTFL clock that, however, are tightly coupled to the TTFL clock. The PTFL mechanisms could take over timekeeping and cause the recovery of behavioral rhythmicity after disruption or disturbance of the TTFL clock. The same posttranslational mechanisms might be at work at the beginning of the knockdown in DD, when circadian locomotor cycle periods continue to increase after mRNA levels are at their minimum. While TTFL clocks were disrupted by the knockdown, underlying redundant timing mechanisms in a system of coupled oscillators appear to be able to reconfigure to a circadian period during the ∼50 days after dsRNA injection, further confirming our systemic hypothesis of circadian timekeeping with our novel modeling. Systemic timing might also occur in other species since similar results have been reported in mice, where *Per2^m/m^* knockouts spontaneously recovered rhythmic locomotor behavior in DD within 20-80 days after arrhythmia onset (Riggle et al., 2022). Reconfiguration and recovery, independent of gene expression, have been also observed in other neural networks with periodic activity (Calabrese and Marder, 2025; Harley et al., 2015; Khorkova and Golowasch, 2007; Luther et al., 2003; More-Potdar and Golowasch, 2023).

The obvious candidate to take over timekeeping duties in cockroach behavior is the network of the circadian clock neurons. Nevertheless, as prerequisite to successful network level synchronization, each individual clock neuron requires endogenous clock mechanisms that compensate for the disrupted transcription-based TTFL circadian clock (Stengl and Schneider, 2024). Accordingly, both *in vivo* and *in vitro* recordings of AMEs in intact brains of the Madeira cockroach show that the prevalence of specific frequencies of ultradian oscillations in the spontaneous neural activity of AME clock neurons express circadian modulation (Rojas et al., 2019; Schneider and Stengl, 2007). In addition to other experimental approaches reported in previous chapters of the discussion, changes in spontaneous electrical activity of AME neurons peak at dusk and dawn, indicating synchronized clock neuron ensemble formation at these ZTs, which requires synchronization through gap junctions (Rojas et al., 2019; Schneider and Stengl, 2007, 2006, 2005). For example, gamma oscillations peaking at dusk are consistent with a synchronized EO clock circuit that releases neuropeptides such as PDF (Rojas et al., 2019; Stengl and Schneider, 2024). Furthermore, cAMP levels, which are upregulated via PDF receptor activation and which modulate pacemaker channel conductances (Lee and MacKinnon, 2017; Pape and McCormick, 1989; Wei et al., 2014), peak at dusk and dawn (Schendzielorz et al., 2014). At these ZTs, maximal changes in electrical activity occur in excised AME clocks, reminiscent of predicted MO and EO activity peaks (Schneider and Stengl, 2007, 2005). Hence, we modeled both the MO and EO as single circadian clock cells that comprises a circadian PTFL membrane clock based on interlinked circadian Ca^2+^ and cAMP oscillations, coupled to a nucleus TTFL clock, based on interlinked circadian *tim1* (or *per*) and *clk* oscillations. Furthermore, depending on the respective light regime, either the MO with short period, or the EO with long period was set to be dominating. Therefore, the results presented here align with our novel hypothesis that the individual circadian clock neurons contain independent but interlinked heterarchical TTFL and PTFL clocks for both robust and flexible timekeeping (Stengl and Schneider, 2024).

### 7.4 Computational modeling supports the view of a heterarchical network

In this study, we have demonstrated that PSAS oscillators are highly adaptable to diverse sets of measured data. The state variables of these oscillators can represent a variety of biological components, such as mRNA levels or Ca^2+^ ion concentrations. In contrast to the common assumption that the TTFL plays a dominant role in shaping circadian behavior, modeling revealed that a heterarchical architecture also appears reasonable for a functioning biological clock. In a hierarchical network, the dominant TTFL suggests a strong leverage from TTFL to PTFL and a much weaker reverse connection and absence of an endogenous PTFL clock. In contrast, the heterarchical network assumed similar functional importance of both TTFL and PTFL clocks, with bidirectional effects of similar strength. Quantifying coupling strengths of oscillators in biological networks is often difficult, as reported in (Schmal et al., 2018). However, by modeling the interconnection by diffusive coupling functions which are linear in state differences combined with time-varying coupling gains, the relationships found in experiments seem to be reflected suitably. The computational model supports the hypothesis that a heterarchical network structure can accurately capture the observed animal behavior: While an increased period in rhythmic activity appeared immediately after “knockdown”, arrhythmic behavior emerged only 14 days after knockdown initiation. Additionally, the increased significance of the PTFL oscillators within the heterarchical network aligned with the recovery of rhythmicity 40 days after knockdown initiation. By adjusting the proposed model to a hierarchical network, simulations failed to reproduce experimental findings. Adjusting the heterarchical DD network to a network in LL allowed us to make predictions that were validated by preliminary biological experiment and need further testing in other species. Our preliminary experiments suggest that the combination of constant light and the knockdown of positive TTFL clock elements lead to an early loss of circadian rhythmicity in locomotor behavior in LL, indicative of interference with the TTFL clock. Raising Madeira cockroaches in LL conditions greatly prolongs post-embryonic development (Barrett and Page, 1989). In addition, LL-raised adult Madeira cockroaches have been reported to express circadian locomotor behavior in DD but not in LL (Page and Barrett, 1989). In our experiments, untreated animals that were raised in LD remained rhythmic in LL for the duration of the recordings, which corresponds to the output of the computational model before knockdown initiation. So far, the only identified cryptochrome in cockroaches is the mammalian-like CRY2 (Werckenthin et al., 2012). Unlike the *Drosophila*-like CRY1, the mammalian-type cryptochromes are transcriptional repressors rather than circadian, light-responsive photoreceptors (Michael et al., 2017). Depending on light, CRY1 activation leads to the degradation of TIM, resetting the circadian clock in *Drosophila* (Busza et al., 2004; Ceriani et al., 1999; Koh et al., 2006; Stanewsky et al., 1998). Our results raise the possibility that Madeira cockroaches also possess CRY1, which would enhance the RNAi-induced disruption of the circadian TTFL in LL conditions: In DKD animals, the positive limb is affected by RNAi, resulting in less *tim* transcription. These low levels of TIM would be highly susceptible to further degradation by constantly activated CRY1. Thus, the combination of constant light and knockdown of the positive limb would affect both positive feedforward and negative feedback pathways of the molecular clock. This possibility may be explored in a revised computational model.

By integrating diverse biological insights into a single mathematical framework, the model complements experimental findings through computational simulations, offering a more comprehensive understanding of circadian regulation. In particular, the model provided insight into how different strengths of coupling between the TTFLs and PTFLs affected the oscillations that modeled the circadian rhythms. The proposed model allows us to predict the behavior of coupled oscillators for different magnitudes of coupling strength much easier as is possible by biological experiments, thus helping in the formulation of new hypotheses on the interconnection between networked neurons and within cells. Thus, we hope that our work will stimulate further testing of our novel hypothesis of systemic timing with coupled TTFL and PTFL circadian clocks in other species.

## 10 Supplementary Material

**Supplementary Figure S 1:**
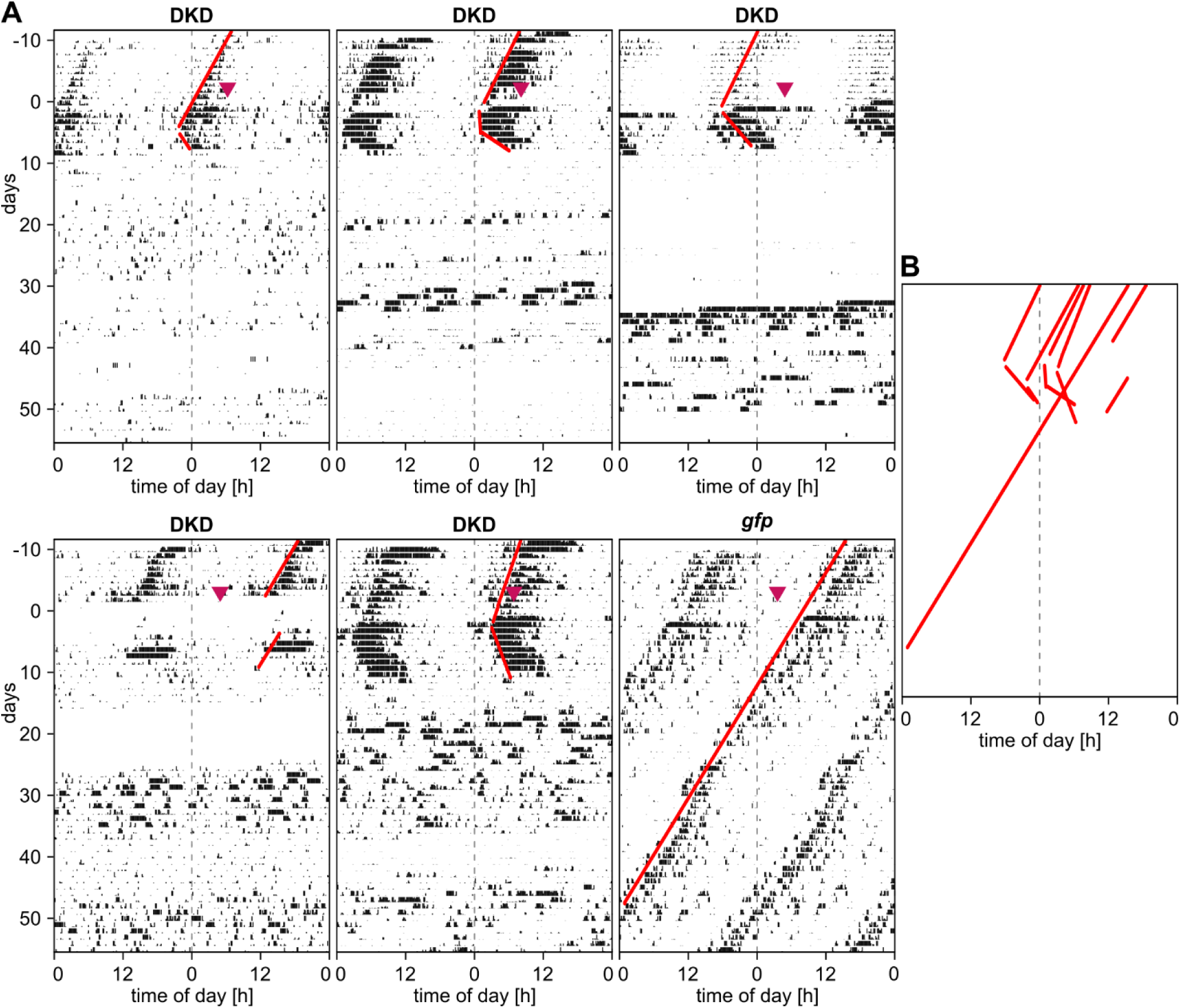
Co-housed cockroaches do not synchronize their locomotor behavior in running wheel assays. **(A)** Double-plotted actograms of six randomly sampled animals housed in the same recording box in the same time window. Each individual was isolated in its own running wheel, but several wheels were co-housed in a recording box, allowing the exchange of chemical cues within one box. Red lines indicate manually assessed onset of circadian activity. Arrowhead indicates the time of double-stranded RNA injection of either *gfp* or double-knockdown (DKD) of *Rm’clk* and *Rm’cyc*. **(B)** Overlay of the onset lines.

**Supplementary Figure S 2:**
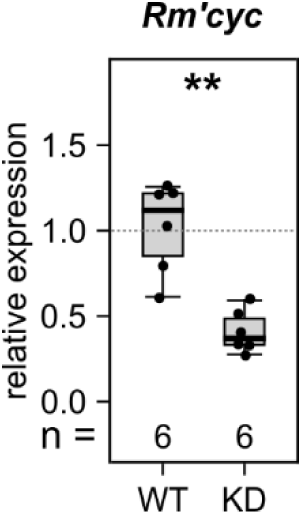
Knockdown efficiency of *Rm’cyc* after 4 weeks in DD. Relative *Rm’cyc* expression was significantly reduced in the knockdown (KD) animals compared to untreated wild-type (WT) controls (Welch’s t-test: t(7.07) = -5.07, p = 0.001).

**Supplementary Figure S 3:**
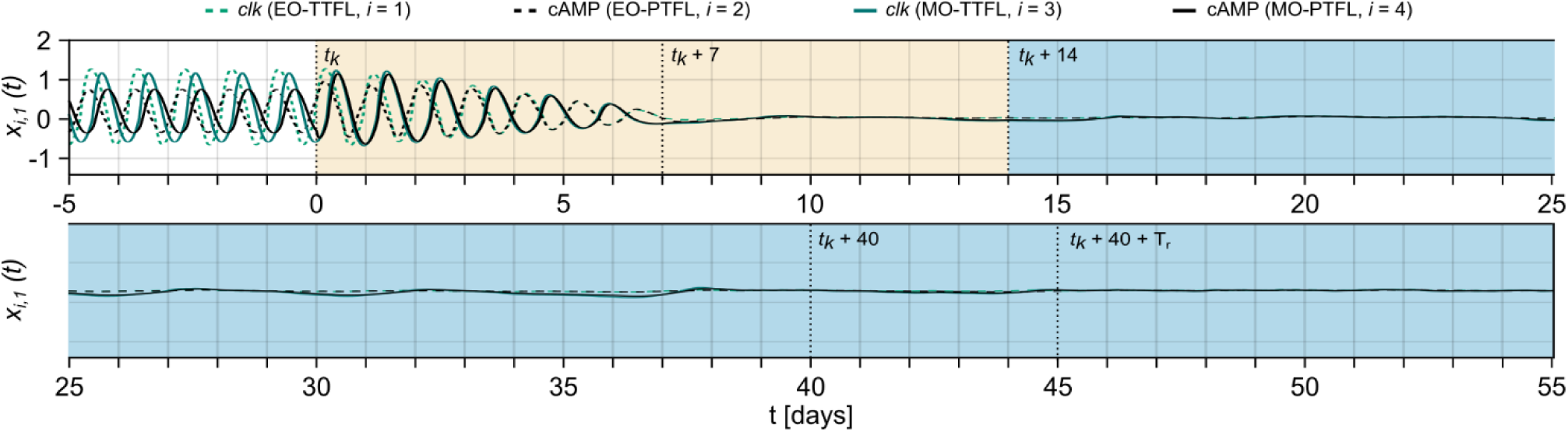
Additional simulation of the hierarchical network for which the *A* matrix of the transient PSAS has real negative eigenvalues. The simulation shows *x_i_*_,1_(*t*) for *i* ∈ {1,2,3,4} for a hierarchical network for which the *A* matrix of the PSAS has real eigenvalues instead of complex eigenvalues (simulation shown in Figure 8). The coupling gains are also adjusted for this case. Relative changes in coupling gains are the same as in Figure 7A.

## 12 Acknowledgements

We thank Denise Rittmeister and Anna-Maria Schrader for helping with the knockdown experiments. We further thank André Arand for rearing the animals, Romy Freund for qPCR assistance and Achim Werckenthin for helpful discussions.

## 13 Author contributions

Conceptualization: OS, MS; Data curation: HZ, OS, MS; Formal analysis: HZ, TT, LK, ACS; Funding acquisition: OS, MS; Investigation: HZ, TT, LK, PP; Methodology: HZ, TT, LK, OS, MS; Project administration: ACS, OS, MS; Resources: OS, MS; Software: HZ, TT, LK, OS; Supervision: HZ, ACS, OS, MS; Validation: HZ, TT, LK, ACS, OS, MS; Visualization: HZ, TT, LK, ACS, MS; Writing – original draft: HZ, TT, LK, ACS, MS; Writing – reviewing C editing: HZ, ACS, OS, MS.

## 14 Statements and declarations

### 14.1 Ethical considerations

Under the applicable regulations, ethics approval was not required for this study, which involved only invertebrates. All work was carried out in accordance with the institutional guidelines at University of Kassel and guidelines of the German Research Foundation (DFG).

### 14.2 Consent to participate

Not applicable.

### 14.3 Consent for publication

Not applicable.

### 14.4 Declaration of conflicting interest

The authors declared no potential conflicts of interest with respect to the research, authorship, and/or publication of this article.

### 14.5 Funding statement

HZ, TT, LK, ACS, OS, and MS were supported in part by Deutsche Forschungsgemeinschaft RTG 2749/1: “Biological Clocks on Multiple Time Scales”.

### 14.6 Data availability statement

The datasets and model generated during and/or analyzed during the current study will be made available in the DaKS data repository of the University of Kassel when the manuscript has been accepted for publication.

## Notes

### Competing Interest Statement

The authors have declared no competing interest.

### Summary of Updates

1)We added two expansions of the model and simulated identical knockdowns as in the original model. One shows the model in a hierarchical configuration in which the TTFL clocks are the only endogenous oscillators (Figure 8, Supplementary Figure S3). The other shows the heterarchical model in constant light (LL) conditions (Figure 9) 2)We tested the prediction of the heterarchical model in LL conditions with cockroaches in running wheel assays (Figure 9). Due to the time limit for the resubmission of the manuscript we were only able to obtain preliminary data, but this data matches the model prediction. 3)We validated knockdown efficiency by qPCR for all 5 genes in our knockdowns (Figure 3D, Supplementary Figure S2), and did new experiments to compare relative mRNA levels in knockdowns and untreated animals of the same age (Figure 6A). 4)We compared circadian activity of the animals housed in the same recording box to rule out synchronization though chemosensory cues (Supplementary Figure S1) 5)We revised Introduction and Discussion

